# Probabilistic Models of Larval Zebrafish Behavior: Structure on Many Scales

**DOI:** 10.1101/672246

**Authors:** Robert Evan Johnson, Scott Linderman, Thomas Panier, Caroline Lei Wee, Erin Song, Kristian Joseph Herrera, Andrew Miller, Florian Engert

## Abstract

Nervous systems have evolved to combine environmental information with internal state to select and generate adaptive behavioral sequences. To better understand these computations and their implementation in neural circuits, natural behavior must be carefully measured and quantified. Here, we collect high spatial resolution video of single zebrafish larvae swimming in a naturalistic environment and develop models of their action selection across exploration and hunting. Zebrafish larvae swim in punctuated bouts separated by longer periods of rest called interbout intervals. We take advantage of this structure by categorizing bouts into discrete types and representing their behavior as labeled sequences of bout-types emitted over time. We then construct probabilistic models – specifically, marked renewal processes – to evaluate how bout-types and interbout intervals are selected by the fish as a function of its internal hunger state, behavioral history, and the locations and properties of nearby prey. Finally, we evaluate the models by their predictive likelihood and their ability to generate realistic trajectories of virtual fish swimming through simulated environments. Our simulations capture multiple timescales of structure in larval zebrafish behavior and expose many ways in which hunger state influences their action selection to promote food seeking during hunger and safety during satiety.

## Introduction

Methods to quantify freely moving animal behavior have seen explosive growth in recent years. This growth has been fueled by rapid improvements in the quality and accessibility of scientific cameras, pose estimation algorithms, and behavioral models. A modern behavioral analysis pipeline^1^ commonly involves: (1) acquiring video of an animal behaving, (2) compressing this data into a low-dimensional time series representing the animal’s posture (or postural dynamics) in each video frame, and (3) using computational tools to annotate each frame with a discrete behavioral state label (e.g. “head-grooming”, “rearing”). While each of these steps pose significant challenges, variations of this pipeline have been used to discover large sets of stereotyped actions generated by worms^2,3,4,5,6^, flies^7,8,9^, fish^10,11,12,13,14^, and mice^15^ behaving in controlled lab environments. Importantly, these procedures produce statistical summaries of behavior that facilitate comparison of nervous system function across animals or animal groups.

The larval zebrafish is an important animal species for investigating the development^16,17,18^, structure^19, 20^, and function^21,22,23^ of vertebrate nervous systems. Further, they are convenient to use in behavioral studies because of their simple body plan, temporally discrete behavior, and stereotyped locomotor repertoire. Pose estimation^24,25,26,27,28^ is an active area of machine learning research and many solutions have been developed to estimate pose from video, even for animals with complicated and flexible bodies such as locusts, fruit flies, and humans. Larval zebrafish, however, have a simpler body shape with fewer degrees of freedom, reducing the complexity of this problem. Once extracted, animal pose data can be temporally segmented into a sequence of minimal and stereotyped behavioral elements (referred to as “movemes”^29^, “actions”^29^, “motifs”^15^, or “syllables”^15^). This parsing problem becomes challenging for animals that move continuously or display many behaviors, but several effective approaches have been described^1^. However, since larval zebrafish swim in punctuated bouts, temporal segmentation of their behavior into bout and interbout epochs is straightforward. Together, the anatomy and movement patterns of larval zebrafish simplify behavioral analyses and allow each swim bout to be represented as a point in a high dimensional posture or kinematic parameter space. Several studies have taken advantage of these properties to analyse and categorize swim bouts, and the most comprehensive^14^ effort to date identified 13 basic types of swims used during hunting^30, 31^, taxis behaviors^32,33,34,35^, escape maneuvers^36,37,38^, social interactions^39^, and spontaneous^40^ swimming in the light and dark.

While locomotor patterns produced by larval zebrafish have been well studied, much less is understood about the complex generative processes underlying the animal’s natural action selection. For example, when a zebrafish larvae swimming through a natural environment selects and generates a swim bout from its locomotor repertoire, what variables and computations shape this decision? How does this decision depend on external inputs like the locations, sizes, shapes, and motion patterns of objects in the local environment? How does this decision depend on internal inputs like short- or long-term behavioural history, or hunger state? To work toward addressing these questions, we use a moving camera system to collect high spatial resolution video of larvae swimming in a large arena with abundant prey. This approach allows the fish to explore and hunt throughout a vast environment as we monitor fine postural features (e.g. tail shape, eye positions) together with microscopic details of its surroundings. Importantly, since zebrafish larvae frequently move their eyes to hunt, stabilize gaze, navigate, and surveil for threats, continuous eye position measurements can provide rich information about the animal’s behavioral state. The use of a moving camera system also eliminates a trade-off between spatial resolution and arena size, allowing us to observe up to hundreds of consecutive swim bouts without interference from arena boundaries. By contrast, fixed camera systems can require small enclosures that physically limit the space of possible behavioral sequences and bias action selection by promoting thigmotactic^41^ behaviors. Here, we avoid these issues to generate behavioral data that should more closely approximate larval zebrafish behavior in the wild. We next use this data to construct probabilistic behavioral models – building on point process models with a long history in computational neuroscience^42,43,44,45^ – to evaluate how internal and external inputs shape larval zebrafish behavior. By sampling from these models, we can simulate behavioral trajectories that capture salient features of larval zebrafish behavioral dynamics spanning multiple timescales, from reactions to prey (*<*100 milliseconds), across stretches of hunting and exploration (seconds to minutes), and throughout changing hunger state (minutes to hours).

## Results

### Acquiring Behavioral Data with BEAST

To collect high resolution video (13 µm per pixel; 60 fps) of larval zebrafish swimming in a large arena, we built a moving camera rig called BEAST (for Behavioral Evaluation Across SpaceTime, **1A**, **Supp Video 1**). This setup is similar to previously described rigs^46^, but the camera moves on a motorized gantry to remain positioned above the fish. We observed 7-8 days-post-fertilization nacre^−/−^ zebrafish larvae (n=130) one at a time. To evaluate the influence of hunger on their behavior, each fish was either given abundant paramecia (*fed* group, n=73) or deprived of food for 2.5-5 hours (*starved* group, n=57) prior to observation. Previous studies have shown that brief food deprivation is sufficient to robustly increase larval zebrafish food intake^47, 48^ and to increase their likelihood to approach, rather than avoid, prey-like visual stimuli^49, 50^. We therefore expect the *fed* and *starved* fish groups to display different behavioral patterns and aim to construct behavioral models to quantify and characterize these effects.

We place each fish in the arena and repeatedly recruit it to the center to initiate up to 18 observational trials. We guide the fish to the center by projecting inwardly drifting concentric gratings onto a screen embedded in the tank bottom, thereby leveraging the fish’s optomotor response^35^. Once at the center, a static natural scene replaces the gratings and the fish is recorded for up to 3 minutes or until it reaches the arena edge or tracking fails. Swim paths from 3 representative trials are shown (**1B**) and the fish’s heading direction throughout the indicated portion (**1B**, purple line) of one trial is plotted (**1C**). Swim bouts can be seen as brief fluctuations in heading direction over time and the timing of bout and interbout epochs is determined from this signal (**1D**, Methods). We translate and rotate each image frame to register the video to the reference frame of the fish, as shown for the indicated 200 millisecond swim bout (**1E**). We encode fish posture (S1) in every image frame by estimating the vergence angle^31^ of each eye and the shape of the tail^46^ (**1F**). Larvae can accelerate rapidly (e.g. during escape swims), which can cause the online tracking system to fail. Also, offline pose estimation is occasionally compromised due to motion blur (during very high speed swims) or body roll (causing one eye to occlude the other in the image). We retain only video segments in which all postural features can be accurately extracted in every image frame for further analysis (see Methods).

The processed dataset contains 40 hours of behavioral data parsed into bout and interbout epochs (4002 video segments) (**1G**). Across all swim bouts (*n* = 200,559), the change in heading angle per bout is narrowly and symmetrically distributed (**1H**). The arena contains abundant prey (mostly paramecia and some rotifers) and the fish tend to hunt prey located near the water surface. Zebrafish larvae converge their eyes during hunts to pursue prey with binocular vision. While not hunting, larvae keep their eyes more diverged, increasing their visual coverage of the environment which should improve their ability to detect threats. These animals should therefore experience a natural trade-off between seeking food and seeking safety, and these opposing states can be seen in the bimodal distribution of eye position measurements during interbout intervals (**1I**). We maintain a large circular field of view centered around the fish head (**1J**) with sufficient image sharpness to extract positions, sizes, shapes, and motion patterns of objects near the water surface (**1K**, Methods) with which the fish may interact. We later use this information to construct compressed representations of environmental state to predict the type of the next swim bout.

### Bout Categorization and Regulation of Action Selection by Hunger State

Hunger state has a large influence on larval zebrafish behavior and this can be seen by comparing eye position histograms (similar to **1I**) across *fed* and *starved* fish groups within the first 10 minutes of testing (**2A**). The fraction of interbout intervals during which eyes are converged (threshold: mean vergence angle = 24°) increased by 183% from 0.124 in *fed* fish to 0.351 in *starved* fish in this time period. *Fed* fish also display wider eye divergence during non-hunting behavior compared to *starved* fish (**2A**, *fed*: peak at 12.5°; *starved*: peak at 15.5°; solid vertical lines). These observations indicate that hunger promotes increased food seeking behavior while satiety promotes wider eye divergence.

To quantify the structure of larval zebrafish behavioral sequences and compare this structure across *fed* and *starved* groups, we first aim to categorize all observed swim bouts into discrete types. To that end, each swim bout is represented as a 10-frame (167 ms) postural sequence beginning at bout initiation (**2B**). This gives a 220-dimensional representation (2 eye and 20 tail measurements per frame) of the postural dynamics associated with each swim bout. We next perform dimensionality reduction to embed these 220-D observations in a 2-D space with t-distributed stochastic neighbor embedding (tSNE^51, 52^) (**2C**) and use density-based clustering^8, 53^ to isolate 5 major classes of swim bouts (S2-S4). These classes consist of hunting bouts (here called *J-turn*, *pursuit*, *abort*, *strike*) and non-hunting bouts (here called *exploratory*). Zebrafish larvae typically initiate hunts by converging their eyes and orienting toward prey with a *J-turn*^31^, and reduce the distance to prey while maintaining eye convergence with *pursuit*^54^ bouts (also called approach^14^ swims). Larvae typically end hunts with eye divergence, either during a *strike* (also called capture^14^ swim) or during hunt termination with an *abort*^55^. We find *starved* fish upregulate use of all hunting bouts, with *strikes* upregulated most and *aborts* least (**2D**). Bout-class selection probabilities of *fed* and *starved* fish gradually converge over ~40 minutes (S5), presumably as the hunger state of each group shifts from their opposing initial conditions (high hunger or high satiety) toward an intermediate state near nutrient equilibrium.

To increase the granularity of the swim bout categories and improve the sensitivity of our analyses, we further subdivide the 3 largest bout-classes to yield a total of 10 exploring bout-types and 8 hunting bout-types (**2E**). Each of these 18 bout-types is composed of nearly equal numbers of leftward and rightward bouts. We use 2 scalar kinematic measurements to subdivide swim bout-classes: |Δheading| and |Δtail-shape| (S6). |Δheading| is the magnitude of heading angle change per bout, and |Δtail-shape| is the sum of the magnitudes of frame-to-frame changes in tail shape across each 10-frame bout representation, a postural measurement that correlates with distance traveled per bout and presumably energy expenditure (see Methods). We evenly split *J-turns* into 2 types (*j1*, *j2*) by |Δheading|. *Pursuits* are more abundant, so we split first by |Δheading| and then again by |Δtail-shape| to yield 4 types (*p1-4*). *Exploratory* bouts are split into 3 groups by |Δheading| and again into 3 groups each by |Δtail-shape| to yield 9 types (*e1-9*). A small subset of *exploratory* bouts (called *e0*) seem to correspond to orofacial and/or pectoral fin movements occuring in the absence of tail motion. *e0* bouts occur with eyes diverged, include actions like suction, jaw movement, swallowing, and prey expulsion, and are the most likely bout-type to follow *strike*. *e0* events were isolated prior to splitting exploratory types *e1-9*. Since *e0* bouts are near threshold for bout detection, we expect this category to also include some erroneously detected bouts due to measurement noise and temporal segmentation errors. Detailed kinematic summaries of all bout-types are found in S6-S7. With labels assigned to all swim bouts, the distributions of interbout intervals preceding and following each of these 36 bout-types can be compared (**2F**, S8). It is apparent that larvae select longer interbout intervals during exploration (i.e. while not hunting), and select shorter interbout intervals during hunts. Note also that distributions of interbout intervals preceding (or following) left and right versions of each bout-type are nearly equivalent, demonstrating the extent to which left-right symmetry organizes larval zebrafish behavior at the population level.

By comparing bout-type abundances (**2G**) and interbout interval durations (**2H**) across *fed* and *starved* fish groups in the first 40 minutes of testing, we find that hunger influences larval zebrafish bout-type selection in previously unreported ways. *Fed* fish upregulate selection of low-energy *exploratory* bouts (*e1-3*) relative to *starved* fish, with selection of the lowest-displacement forward swim *e1* increased by 107%. *Starved* fish are more likely to start hunts, especially through high-angle *J-turns* (*j1* up 35%, *j2* up 70%), and increase use of *pursuits* (up 71%), *aborts* (up 34%), and *strikes* (up 96%). Hunger state also affects interbout interval selection, especially following *exploratory* bouts, with *fed* fish selecting longer interbout intervals. This effect is most pronounced following low-energy *exploratory* bouts, with the mean duration of interbout intervals following bout-types *e1-3* increased by 101 ms in *fed* fish relative to *starved* fish. Also, during interbout intervals preceding exploring bouts, *fed* fish maintain wider eye divergence than *starved* fish. (**2I**). Taken together, these results indicate that satiation promotes multiple strategies that should increase protection against predation. *Fed* fish might decrease their visibility to predators by upregulating low-energy *exploratory* bouts and spending more time at rest (i.e. longer interbout intervals). *Fed* fish might also be better able to detect predators by maintaining wider eye divergence while exploring (low-energy *exploratory* bouts have highest eye divergence, see S7). By contrast, *starved* fish swim more frequently and cover more distance per bout while exploring. *Starved* fish should also increase their food intake by starting more hunts and increasing the likelihood that hunts end with *strike*.

In these experiments, zebrafish larvae alternate between modes of exploring and hunting, with each mode defined by distinct behavioral dynamics. This mode-switching is highlighted in an example bout sequence containing a successful hunt (**2J**, **Supplementary Video 2**). The trajectory of the fish throughout this hunt is reconstructed (**2K**) and plotted together with 999 other hunts that also end in *strike* (**2L**). We define a complete hunt as a bout sequence that begins with a *J-turn*, ends with an *abort* or *strike*, and is padded with only *pursuits* (for hunts longer than 2 bouts). The full dataset contains 7230 complete hunts (19.6% end in *strike*). To better quantify the behavioral sequences observed in this study, we next construct probabilistic models to predict the timing and type of swim bouts.

### Constructing Probabilistic Models to Predict Interbout Intervals

We model the data as a marked renewal process^45, 56^, a stochastic process that generates a sequence of discrete events in time, each characterized by an associated “mark” (**3A**). Marked renewal processes are statistical models that specify the conditional distribution of the time and type of the next event in a sequence given the history of preceding events. First, we consider bout timing. Our key question for model construction is, “what features of the event history carry predictive information about the timing of the next event?” To this end, we choose five interpretable features to represent behavioral history across multiple timescales (**3B**). On the shortest timescale, we model the interbout interval (*i_n_*) as a function of *preceding bout-type* (*b_n−_*_1_) and *preceding interbout interval* (*i_n−_*_1_). On an intermediate timescale, we use *hunt dwell-time* (*t*_hunt_) and *explore dwell-time* (*t*_explore_) features to encode how long the fish has been dwelling in either a hunting or exploring mode just prior to *i_n_* (see Methods). On the longest timescale, we encode how long the fish has been in the tank prior to *i_n_*(*tank-time*, *t*_tank_), which can relate how fish behavior changes with hunger state. By comparing models composed from different features, we can learn how past actions predict future behavior.

We found that *starved* fish select shorter interbout intervals than *fed* fish, but how else do patterns of bout timing differ across fish groups? To interpret how feeding state influences the functional relationship between behavioral history and interbout interval selection, we consider 2 forms for each predictive feature: *pooled* and *split*. In the *pooled* form, data from *fed* and *starved* fish are pooled together to fit one set of weights relating that predictive feature to interbout *i_n_*. In the *split* form, a separate set of feature weights are fit for each fish group. We use a generalized linear model^57^ (GLM) with an exponential inverse link function to generate a probability distribution over *i_n_*(**3C**, Technical Appendix). Briefly, the dot product of a basis function representation (S9) of the feature input with the corresponding feature weights is computed. This value is passed through an inverse link function to give the mean of a probability distribution over *i_n_*. Since the interbout intervals are measured in units of frames elapsed, we considered 3 types of probability distributions over nonnegative counts: *geometric*, *Poisson*, and *negative binomial* (*NB*). We find the data to be too complex for modeling with *geometric* or *Poisson* distributions, which are parameterized by just their mean. Instead, we can fit the data better with the more flexible *NB* distribution, which is parameterized by its mean and variance. This requires an additional set of feature weights to estimate the variance of the *i_n_* distribution.

In **3D-F**, we visualize results from fitting *pooled* and *split* forms of models composed from each intrinsic feature listed in **3B**. For each model, we select an appropriate prior variance on the weights through an empirical Bayes^58^ hyperparameter selection method (Technical Appendix), and for features that take a scalar value as input (those in **3F**), we also choose the number of basis functions. We compute the marginal log likelihood (MLL) of each model to select hyper-parameters and to choose a preferred feature form (*pooled* or *split*, indicated with gold star). In modeling the interbout interval as a function of *preceding bout-type*, a separate *i_n_* distribution is produced for each of the 18 possible values for *b_n−_*_1_ (with 2 example values shown, **3D**). The *NB* distribution fits observed data better than *geometric* or *Poisson*, so we use the *NB* throughout the rest of the paper. With a *split* form, the *preceding bout-type* feature captures subtle bout-type specific differences in interbout interval durations across *fed* and *starved* fish groups (**3E**). For each remaining feature, we report the predictive mean of the *NB* distribution for *fed* and *starved* groups across a range of possible input values (**3F**). We include an additional set of *split-bias* weights in each model in **3F** to capture the overall mean and variance of interbout intervals for each fish group (2 extra free parameters per fish group). This design choice allows models composed from *pooled* features (**3F**, top row) to capture shifts in mean and variance of interbout intervals across fish groups without using the more complex *split* features (**3F**, bottom row).

After fitting the interbout models, we next visualize their predictive output to see how behavioral history influences future interbout interval durations selected by the fish. On a short timescale, we find consecutive interbout interval durations are autocorrelated (**3F**, column 1). On an intermediate timescale, interbout intervals get shorter as hunt sequences get longer (**3F**, column 2). By contrast, as exploring sequences get longer, interbout intervals also get longer (**3F**, column 3). After accounting for the shift in mean, the relationship between these 3 features and interval *i_n_* is similar across fish groups. By contrast, *fed* and *starved* fish display opposing patterns of interbout interval selection on the longest timescale (**3F**, column 4). *Starved* fish initially hunt more, producing shorter interbout intervals. *Fed* fish initially explore more, producing longer interbout intervals. As their hunger states equilibrate, interbout intervals selected by *fed* and *starved* fish become more similar. These opposing behavioral patterns require the *split* form of the *t*_tank_ feature to be modeled appropriately.

### Constructing Probabilistic Models to Predict Bout-Types

The second component of the marked renewal process is a model of how the next bout-type is selected depending on behavioral history, including the interval immediately preceding it. We use the previously introduced intrinsic features (from Figure 3) and include also 4 extrinsic features (**4A**) to model how locations (*v*_loc_), sizes (*v*_size_), and relative velocities (*v_x_*, *v_y_*) of objects in the fish’s local environment relate to bout-type selection. We use locations of putative prey objects to construct an 868-dimensional image (*v*_loc_) encoding their final positions prior to initiation of the next swim bout. We modify *v*_loc_ to produce the other extrinsic features by scaling the intensity of each represented object (S9), with an example *v*_size_ input shown (**4A**). Similar to the interbout models, we take the dot product of a basis function representation of a feature input with its corresponding weights to produce a vector of bout-type “activations”, *ψ* (**4B**). This vector *ψ* is passed through a softmax function to generate a valid probability distribution, *π*, over all 36 possible bout-types (Technical Appendix). As before, we perform hyper-parameter selection for all features (S10) and select a preferred form (*pooled* or *split*). We also include a set of *split-bias* weights to account for differences in baseline bout-type abundances across *fed* and *starved* fish groups (36 extra free parameters per fish group). This again allows simpler *pooled* models to account for differences in bout-type abundances across groups, and we choose *pooled* forms for all bout-predicting features except *t*_tank_.

**Figure 1:**
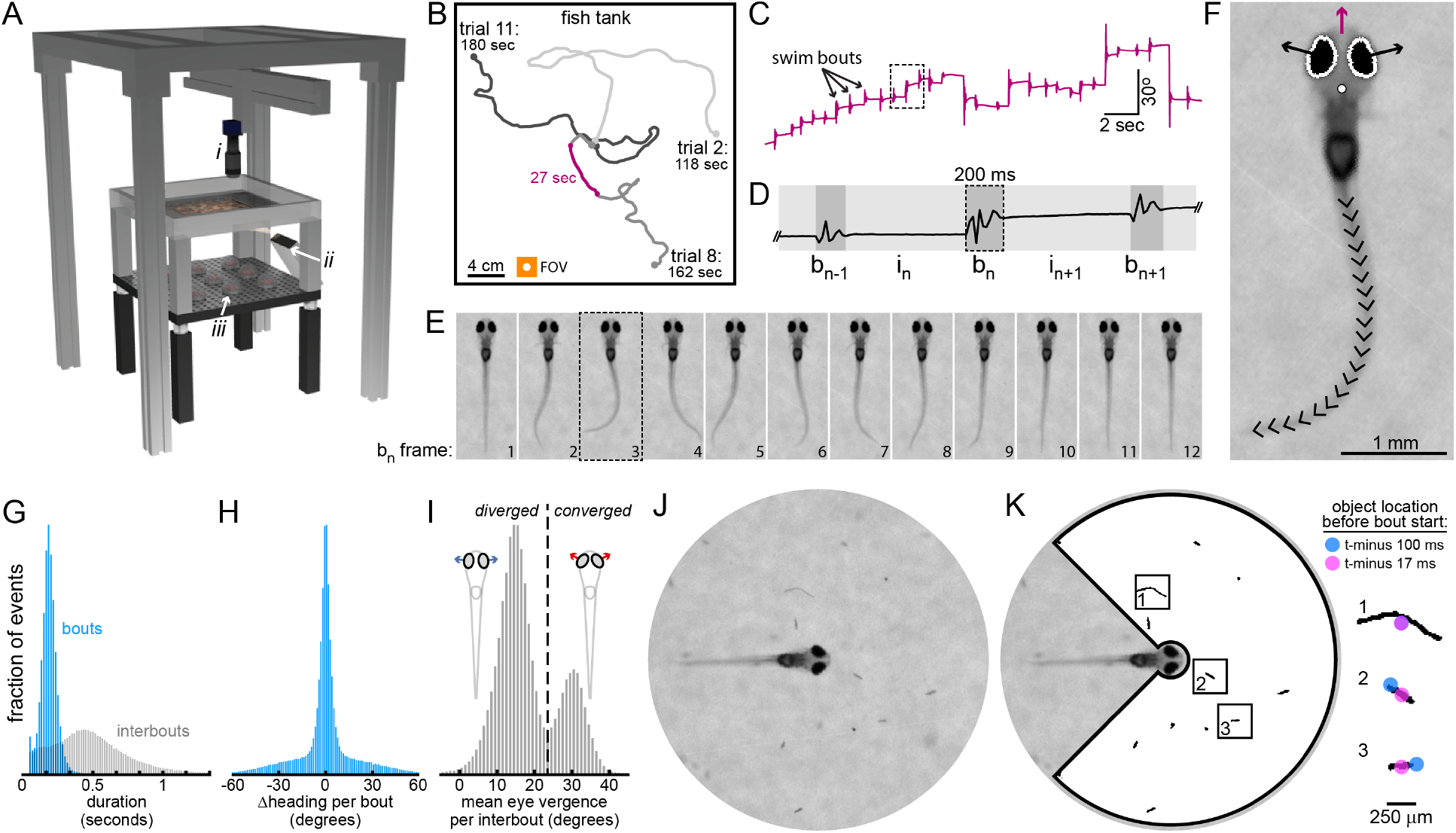
Acquiring larval zebrafish behavioral data with BEAST. **A.** Schematic of behavioral rig. An infrared camera (i) moves on a motorized gantry to remain centered above a freely swimming fish. A projector (ii) and IR-LEDs (iii) illuminate a diffusive screen embedded in tank bottom. An air table isolates the fish from vibration (see **Supplementary Video 1**). **B.** Swim paths from 3 trials from 1 fish are shown. Fish tank dimensions = 300 × 300 × 4 mm. The size of the total camera field of view (FOV) is shown (orange square: side length = 22 mm) **C.** Fish heading direction during 27-second excerpt from **B** is shown. **D.** Fish heading direction during 2-second excerpt from **C** is shown. Timing of bout and interbout epochs is determined from changes in heading direction over time (see Methods), and notation for behavioral sequences is indicated. **E.** Each image frame from 200 ms swim bout from **D** is aligned to maintain head position and heading direction. **F.** Fish head position (white circle), heading direction (purple arrow), eye vergence angles (angles between black arrows and horizontal), and tail shape (20 local tail tangent angles indicated with black carets) are extracted in every image frame (shown here for frame indicated in **E**). See S1 for posture estimation details. **G.** Histograms of bout and interbout durations from whole dataset. **H.** Histogram of heading direction change per bout (Δheading) for all bouts. **I.** Histogram of mean eye vergence per interbout for all interbouts. For each interbout, all eye vergence measurements are averaged. Larvae typically explore with eyes diverged and hunt with eyes converged. Local minimum at 24°(dashed line). **J.** The field of view in the video dataset is large enough to see the fish and its local environment (circle diameter = 8.12 mm, size indicated as circle within orange square in **B**). The focal plane is near the water surface. **K.** Locations, sizes, shapes, and relative velocities of all found surface objects preceding swim bout initiation are extracted. Relative motion of 3 objects are shown. In the next image frame, the fish initiates a swim bout (shown in **2B**) to pursue the prey object in **box 2**. Found objects are mostly edible (paramecia, rotifers) but include also algae, dust (like in **box 1**), and some artefacts. Box width = 820 µm.

**Figure 2:**
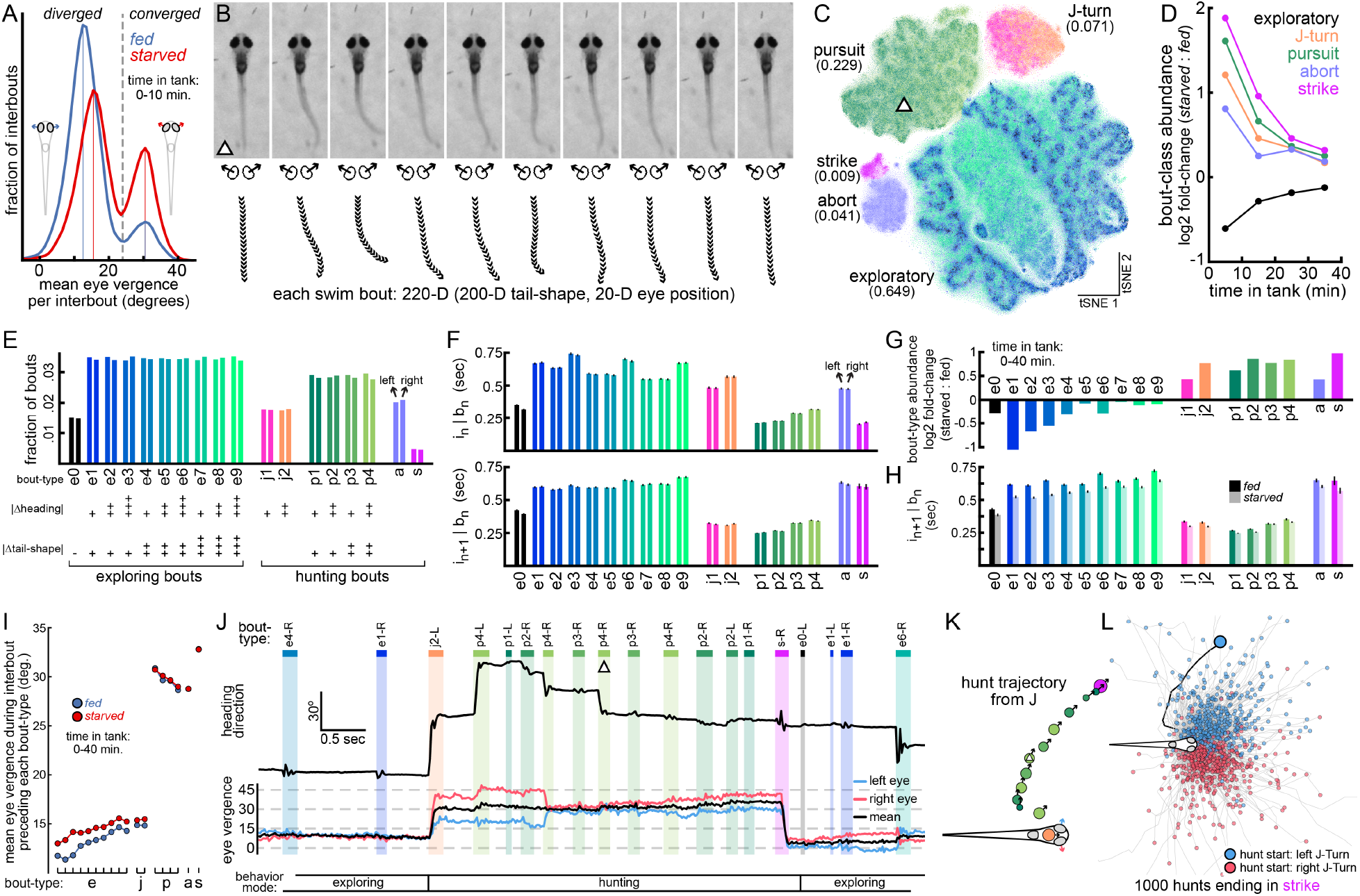
Bout categorization and regulation of action selection by hunger state. **A.** Histogram of mean eye vergence per interbout (as in **1I**) for interbouts observed in first 10 minutes of testing, plotted separately for *fed* and *starved* fish groups. Dashed vertical line indicates threshold between divergence and convergence. Solid vertical lines indicate local histogram peaks. **B.** Each swim bout is encoded as a sequence of postures through 10 consecutive image frames (167 ms) beginning at bout initiation. **C.** The 220-D bout dataset is reduced to 2-D with tSNE and 5 bout-classes are extracted with density-based clustering (S2). Bout-class abundances are shown in parentheses. The location in tSNE space of the swim bout from **B** is indicated with a triangle. **D.** The log_2_ of relative bout-class abundances (*starved* / *fed*) is calculated in 10-minute time bins. **E.** *Exploratory*, *J-turn*, and *pursuit* bout-classes are subdivided by |Δheading|. *Exploratory* and *pursuit* bouts are further subdivided by |Δtail-shape|. Plus signs indicate kinematic parameter magnitude (see S6 for subdivision details). **F.** Durations (mean SE) of interbout intervals preceding (top panel) and following (bottom panel) left and right versions of each bout-type. **G.** The log_2_ of relative bout-type abundances (*starved* / *fed*) is calculated for all bouts observed in first 40 minutes of testing. **H.** Durations (mean SE) of interbout intervals following each bout-type for all bouts in first 40 minutes of testing are shown for *fed* (darker bars) and *starved* (lighter bars) fish groups. **I.** Mean eye vergence during interbout interval preceding each bout-type across *fed* and *starved* fish (time in tank *<* 40 min). Note that hunger state affects eye vergence preceding *exploratory* bouts and *J-turns*, but not preceding *pursuits*, *aborts*, and *strikes*. **J.** Example behavioral sequence in which the fish exits exploring mode to execute a 13-bout hunt ending with *strike*, after which it returns to exploring. Fish heading direction and eye vergence angles are shown with bout and interbout epochs indicated. Bout from **B** indicated with triangle. **K.** Trajectory of hunt from **J**. Cartoon fish length = 4 mm. Circle locations and arrow directions indicate fish head location and heading direction prior to initiation of each bout. Circle color indicates bout-type. Circle area is proportional to |Δtail-shape*|* for each bout. **L.** Trajectories from 1000 randomly selected complete hunts ending in *strike*.

**Figure 3:**
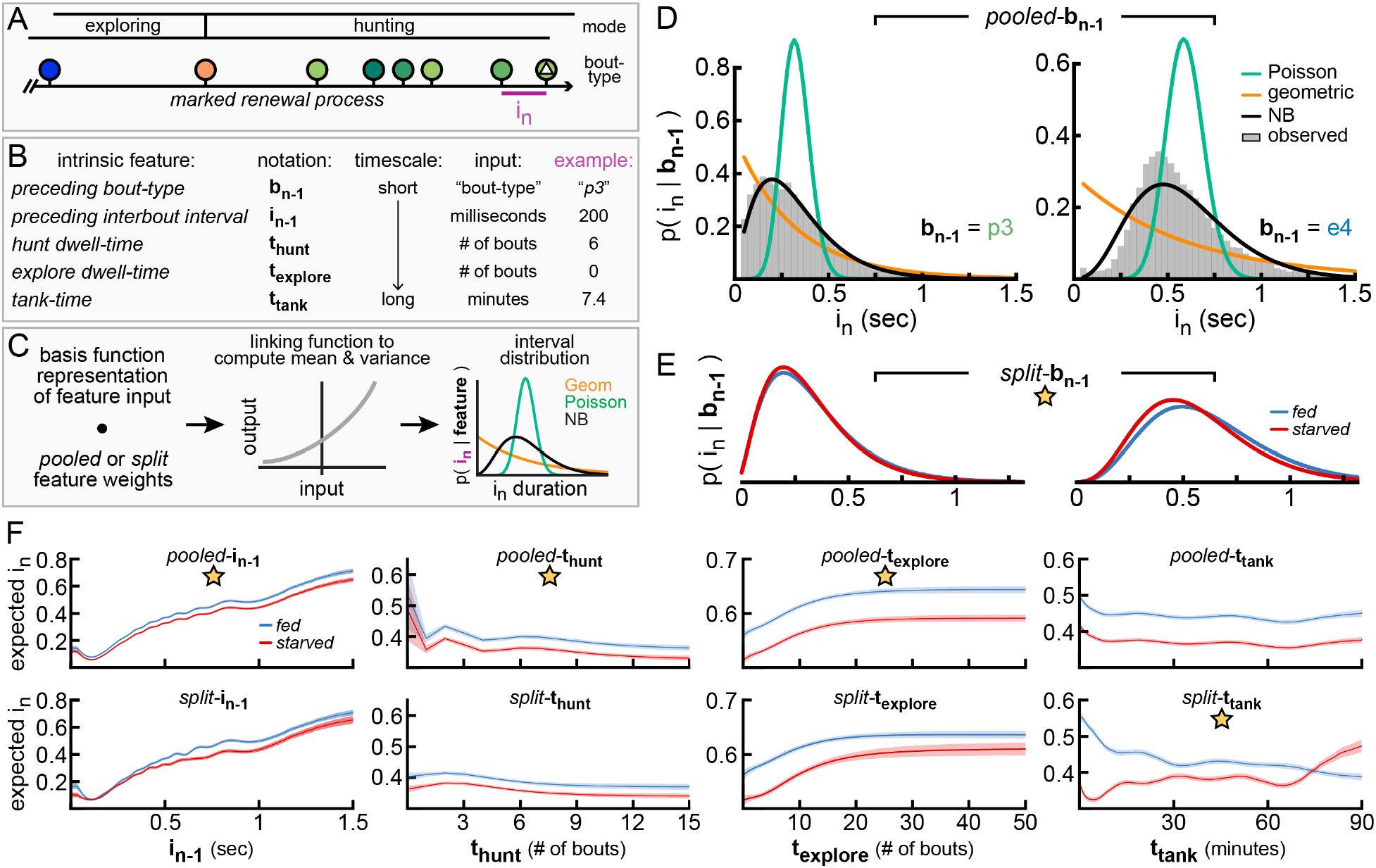
Probabilistic models to predict interbout interval duration. **A.** Behavioral data is conceptualized as a marked renewal process, here for a portion of the sequence shown in **2I**. **B.** Five intrinsic features to model *i_n_*. Example input values are shown to predict duration of *i_n_* indicated in **A**. *Preceding bout-type* is the bout-type assigned to *b_n_*_−1_ and is a categorical feature encoded with a one-hot indicator. Remaining features are temporal features encoded with radial basis functions (see Methods and S9). *Preceding interbout interval* is the duration of *i_n_*_−1_. *Hunt dwell-time* is the integer number of observed preceding consecutive hunting bouts. *Explore dwell-time* is the integer number of observed preceding consecutive exploring bouts. Either *t*_hunt_ or *t*_explore_ is equal to zero for each *i_n_*. *Tank-time* is the time elapsed since the first trial was initiated for that fish. **C.** Schematic of GLM to generate a probability distribution over interbout interval values as a prediction for *i_n_*, measured in units of elapsed frames. For each feature, both a *pooled* and *split* form are constructed. For the *pooled* form, a single set of feature weights is fit for all fish, pooling data across *fed* and *starved* groups. For the *split* form, separate sets of feature weights are fit for each fish group. Either the *pooled* or *split* form is selected based on marginal log likelihood. See Technical Appendix. **D.** For 2 example values of *preceding bout-type* (*b_n_*_−1_), distributions over *i_n_* are generated from *pooled*-*b_n_*_1_ models. The negative binomial (*NB*) distribution fits observed interbout interval distributions best. **E.** For the same example *b_n_*_−1_ values as **D**, separate distributions for *fed* and *starve*d fish are generated from the *split*-*b_n_*_−1_ *NB* model. For this feature, the *split* form is preferred, indicated with gold star. **F.** Predicted *i_n_* values (mean and 95% posterior credible intervals) for *fed* and *starved* fish from *NB* models which take as input *preceding interbout interval* (column 1), *hunt dwell-time* (column 2), *explore dwell-time* (column 3), and *tank-time* (column 4). *Pooled* (top row) and *split* (bottom row) feature forms are shown, with preferred form indicated. All 8 models here include 4 extra parameters (*split-bias* weights) to separately capture mean and variance of *fed* and *starved* interbout intervals.

The tendency of larval zebrafish to switch between modes of exploring and hunting can be seen in the block structure of bout transition probabilities captured by the *preceding bout-type* feature (**4C-D**). While exploring, larvae are likely to link consecutive *exploratory* bouts of similar energy (note increased transition probability along diagonal in **4C**). This bout transition pattern may permit larvae to maintain speed while exploring, especially in combination with autocorrelated interbout intervals. It is known that larval zebrafish can adjust their swimming speed to match optic flow stimuli^59^, and it seems they employ speed control mechanisms during natural exploratory behavior as well. Larvae enter hunting mode with a transition to *J-turn*, after which they are likely to emit some *pursuit* bouts before an *abort* or *strike*. Larval zebrafish behavior is symmetric at the population level, so we construct symmetric features to simplify and improve models. We exploit symmetry in the *preceding bout-type* feature by considering ipsilateral (**4C**) and contralateral (**4D**) bout transition probabilities. It has been shown that zebrafish larvae tend to link bouts in the same direction while exploring a featureless environment, and that neural circuits in the anterior rhombencephalic turning region (ARTR) mediate switching between leftward and rightward states^40^. We too see that larvae tend to link exploring bouts through ipsilateral bout transitions (**4C**) rather than contralateral transitions (**4D**). This pattern extends also to hunting, with nearly every transition being more likely to occur ipsilaterally, except for transitions into *abort* (**4D**, arrow), during which fish are likely to switch left-right state. While bout transition probabilities are different across *fed* and *starved* fish, the transition structure is similar enough that the conservative model selection approach favors the simpler *pooled*-*b_n−_*_1_ form (with 648 feature weights) over the *split*-*b_n−_*_1_ form (with 1296 feature weights). For comparison, see the *split*-*b_n−_*_1_ model (S11).

Prey object locations also influence bout-type selection, especially during hunts, and *pooled*-*v*_loc_ weights associated with each hunting bout-type are shown (**4E**). Since *v*_loc_ inputs are high-dimensional, we compress them by summing *v*_loc_ pixels within spatial bins (as shown). Objects located in red spatial bins increase the probability that the corresponding bout-type will be selected next by the fish. Larvae typically select a *J-turn* to orient toward a laterally located object (~15-60° relative to heading), and the magnitude of heading angle change depends on prey location^30^ (compare *j1*, *j2*). Energetic *pursuits* (*p3-4*) are more likely when prey are further away, while larger |Δheading| *pursuits* (*p2*, *p4*) are more likely when prey are located more laterally. Prey locations also influence how hunts will end, with *strike* becoming most likely with an object located almost directly in front of the mouth. By contrast, *aborts* are only weakly related to prey location and may be selected by the fish to terminate unsuccessful hunts. The *v*_loc_ weights associated with exploring bouts are more spatially uniform with near-zero or negative weights (not shown), increasing the likelihood of exploring bout selection when prey are absent. Taken together, prey locations relate to larval zebrafish bout-type selection in expected ways, and these basic relationships are captured by the *pooled*-*v*_loc_ feature.

The generative process underlying bout-type selection depends nonlinearly on the *preceding interbout interval*, and we capture this dependency with the *pooled*-*i_n_* model (**4F**). The activations (*ψ*) of 5 separate bout-types across the range of *i_n_* input values are shown (**4F**, left panel), with larger activations indicating higher bout-type probability. Full bout-type probability distributions (*π*) evaluated at 3 example values of *i_n_* are also shown (**4F**, panels i-iii), with probabilities across *fed* and *starved* fish averaged for display. At very short *preceding interbout interval* values (*i_n_ <* 0.25 sec), activations of *p3*, *strike*, and *abort* have different dynamics, indicating how timing is intricately involved in bout-type generation during hunts. As *i_n_* extends from 0 to 0.25 seconds, *strikes* become less likely, *aborts* become more likely, and *p3* probability peaks near 150 ms before decreasing again. As *i_n_* reaches 0.5 seconds (**4F**, panel ii), exploring bouts become more probable. As *i_n_* reaches 1 second (**4F**, panel iii), low-energy *e1-3* and high-|Δheading*| e3*, *e6*, *e9* bouts become most likely.

Longer timescale bout-type dependencies are captured with *t*_hunt_ and *t*_explore_ features (**4G-H**). As hunts extend, *aborts* become less likely and *strikes* become more likely (**4G**, left panel). *Pursuits* are always likely when larvae are in hunting mode (i.e. *t*_hunt_ is non-zero), but the probability mass shifts toward the short straight *p1* bout as hunts get longer (**4G**, panel iii) and larvae have approached a target. In these experiments, larvae spend most of their time exploring, and bout-type probabilities shift slightly as larvae spend more time in exploring mode. As *explore dwell-time* increases, larvae become more likely to select the slow straight *e1* bout (**4H**) and also become more locked into the exploring behavioral state. The probability of emitting a *J-turn* decreases ~46% as *t*_explore_ increases from 5 bouts (**4H**, panel i) to 40 bouts (**4H**, panel iii). This effect may be partially explained by decreased food density as fish navigate toward the arena edge. On the longest timescale, the *t*_tank_ feature captures slow fluctuations in bout-type probabilities over the course of observation. As with the models of bout timing, the *t*_tank_ feature must be *split* to capture opposing behavioral trends of *fed* and *starved* fish on this timescale. Separate bout-type probability distributions for *fed* and *starved* fish are shown (**4I**, panels i-iii) and bout-type selection becomes similar across fish groups as their hunger states converge at *t*_tank_ = 40 minutes (**4I**, panel iii).

### Comparing and Combining Behavioral Models

Having constructed several single-feature models of larval zebrafish behavior, we next aim to compare their quality. To this end, we compute the marginal log likelihood (MLL) of each model and report improvement over the simplest baseline model (*pooled-bias*). For interbout models (**5A**), the baseline *NB pooled-bias* model has 2 parameters (1 for mean, 1 for variance). For model comparison, we show MLL for *pooled* and *split* forms of every model introduced in Figure 3. We include also *b_n−_*_2_ as well as *i_n−_*_2_, *i_n−_*_3_, *i_n−_*_4_, and *i_n−_*_5_ to see how predictive information decays over time. We find that *preceding bout-type* is best able to predict *i_n_*, but that *preceding interbout interval* is a close second, and that more distant behavioral history also provides useful information to model the interbout interval. In general, MLL is similar across *pooled* and *split* forms for each feature, except for *t*_tank_. We interpret this to suggest that any mild gains in predictive performance through use of *split* features may be offset by their increased complexity and decreased number of training examples per *split* weight. We therefore select the *pooled* form for all features except those indicated with an asterisk.

While *preceding bout-type* is the best predictor of *i_n_*, can we build stronger models by combining features? More generally, to what extent do the representations of behavioral history encode non-redundant information about interbout *i_n_*? We approach these questions by combining all pairs of features (in their selected forms) to produce 45 paired interbout GLMs (**5B**). For each paired model, we compute MLL (as in **5A**) and report improvement over the stronger of its 2 feature components. Paired models that show large improvement should combine features that provide some unique information about interbout *i_n_*to improve model accuracy. We find that *preceding bout-type* and *preceding interbout interval* combine to produce the strongest paired interbout model (white circle, **5B**). This paired [*b_n−_*_1_, *i_n−_*_1_] model improves over *b_n−_*_1_ alone by 0.05 nats per interbout, or 29% relative to baseline. By contrast, the paired [*b_n−_*_1_, *t*_explore_] model (white square, **5B**) improves over *b_n−_*_1_ by just 0.002 nats per interbout, or 1.4%. Since the models can improve by combining features, we combine all features from **5A** to construct a *combo* interbout model (**5C**). We add features one at a time via greedy stepwise selection, adding the feature that increases MLL most at each step. Model quality improves and eventually saturates during *combo* model construction (note plateau in MLL).

We next repeat this procedure for the bout-type models (**5D-F**). We include also results from GLMs composed from extra extrinsic features listed in **4A** (*v*_size_, *v_x_*, and *v_y_*). We find that *preceding bout-type* is by far the best predictor of bout-type *b_n_*, followed by *hunt dwell-time* and *explore dwell-time*, and then *preceding interbout interval*. As constructed, the GLMs fail to fully capture the complex relationship between the state of the fish’s local environment and its bout-type selection. This problem is challenging for several reasons. Identified environmental objects include both prey (e.g. paramecia, rotifers) and non-prey (e.g. dust, algae), which differentially influence larval zebrafish behavior. Second, environmental objects are abundant in these experiments (mean # of identified objects per bout = 12). This complicates visual scenes encountered by the fish and also our representations of environmental state. Third, the locations, sizes, shapes, and motion patterns of nearby objects are likely to interact in complex ways to influence larval zebrafish action selection. To improve our understanding of how sensory input relates to bout-type selection, we construct feed-forward neural networks that take the extrinsic features as input (S12) and combine them nonlinearly to form a prediction. We find this neural network model improves substantially over the *v*_loc_ GLM (see S12), with predictions of all bout-types improving on held-out data, especially hunting bouts. However, bout-type *b_n_* is still far better predicted by *preceding bout-type*. This result indicates that more sophisticated modeling approaches may be necessary to more effectively predict future larval zebrafish behavior from naturalistic environmental data, but also that bout-type selection depends strongly on the animal’s very short-term behavioral history^1^.

The strongest paired bout model (white circle, **5E**) again combines *preceding bout-type* with *preceding interbout interval*, even though *preceding interbout interval* is just the 4^th^ strongest individual feature. This paired [*b_n−_*_1_, *i_n_*] model improves over *b_n−_*_1_ alone by 0.09 nats per bout, or 14%. By contrast, the paired [*b_n−_*_1_, *t*_explore_] model (white square, **5E**) improves over *b_n−_*_1_ by just 0.005 nats per bout, or 0.8%. This result again indicates that distinct features of the fish’s short-term behavioral history (i.e. *preceding bout-type* and *preceding interbout interval*) encode non-redundant information about its future action selection. As before, we construct a *combo* bout model by iteratively adding features (**5F**). The strongest 3-component bout model ([*b_n−_*_1_, *i_n_*, *v*_loc_]) adds information about prey locations to the strongest paired bout model, combining sources of internal and external information over a timescale of a second or less. Model quality again saturates during construction of the *combo* bout model.

We next take a closer look at the quality of the *combo* interbout and bout models. To confirm the *NB* distribution was a proper choice, we compare the *NB combo* interbout model (from **5C**) to similar *combo* interbout models constructed instead with either *Poisson* or *geometric* distributions (**5G**). While it requires more free parameters, the *NB* model clearly outperforms the others on held-out data. For the bout models, we have so far considered overall quality by assessing how well bout-types can be predicted in general. Here we show predictive performance of the *combo* bout model for each individual bout-type (**5H**), as measured by the F_1_ score^60^ of each one-vs-rest classifier. We compare this performance to that of the baseline *pooled-bias* bout model (**5H**, white lines). This shows us which bout-types are easiest to predict (e.g. *pursuits*, *aborts*, *strikes*, *e0*), and which are most challenging (e.g. *e4-6*, *j2*). In addition, we reproduce this analysis for each of the single feature bout models described in Figure 4, and also for the strongest paired and 3-component bout models (S13). We find the *combo* bout model distributes probability mass over similar bout-types (**5I**), which should be expected if the generative processes involved in the production of similar bout-types are also similar.

**Figure 4:**
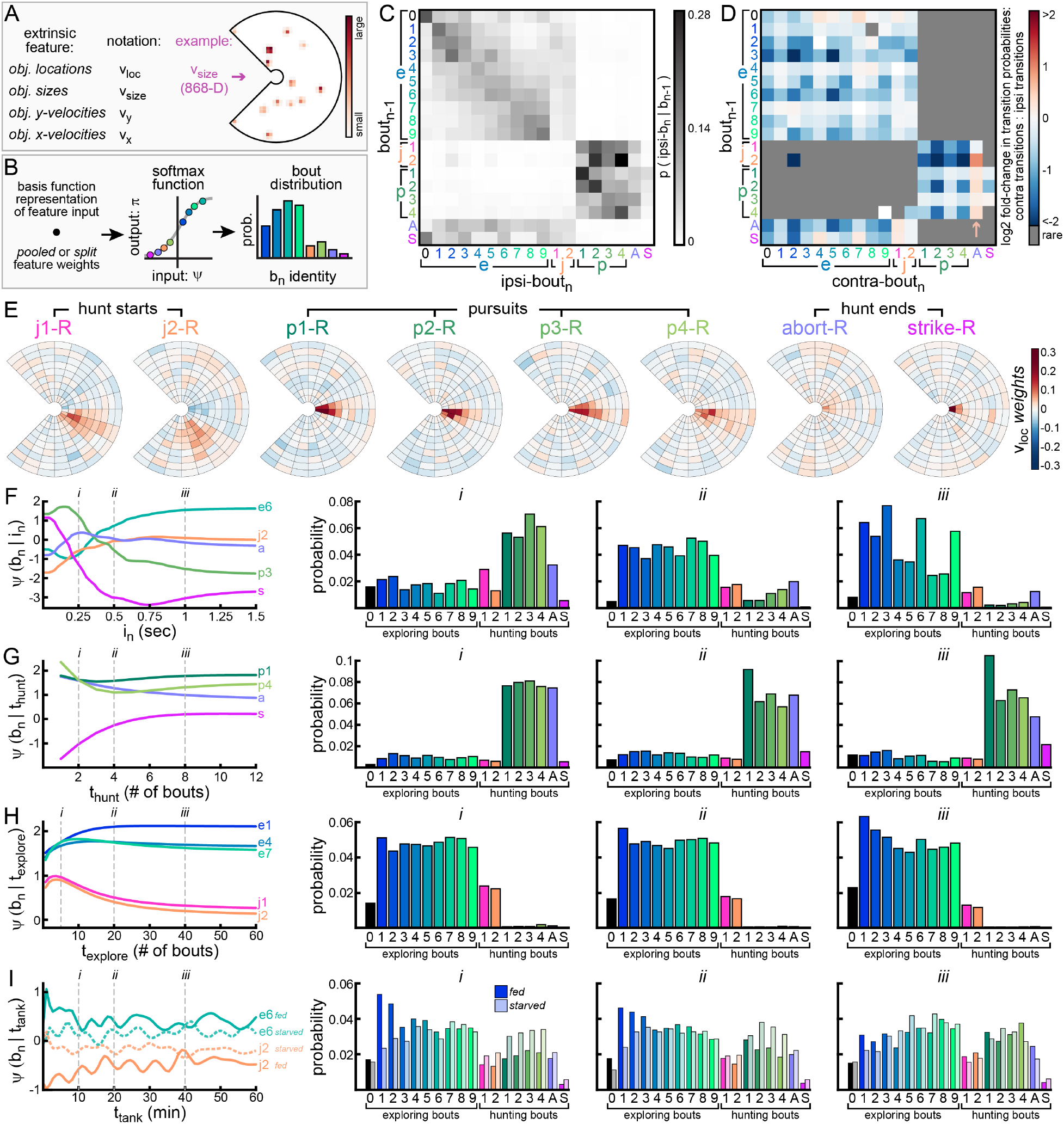
Probabilistic models to predict bout-type. **A.** Four extrinsic features encode locations and properties of found surface objects. See S9 for details. Here, the sizes of objects found in **1K** are encoded to predict the identity of the next swim bout (the *p4-R* bout shown in **2B**). **B.** Schematic of GLM to compute bout-type “activations” (*ψ*) which pass through a softmax function to generate a probability distribution (*π*) over bout-types as a prediction for bout *b_n_*. See Technical Appendix. **C.** Ipsilateral bout transition probabilities (i.e. left to left, or right to right) from *pooled*-*b_n_*_−1_ model. Each row shows the probability that *b_n_* will be each bout-type, given the *preceding bout-type*. **D.** Contralateral bout transition probabilities (i.e. left to right, or right to left) from *pooled*-*b_n_*_−1_ model, reported relative to ipsilateral transition probabilities. Rare transitions are not shown (total transition probability of *b_n_*_−1_ *b_n_ <* 0.015). Blue squares indicate contralateral transitions that are less likely than corresponding ipsilateral transitions. Nearly all transitions are more likely to be ipsilateral, except from *j2* and *p1-4* into *abort* (arrow). **E.** Weights from *pooled*-*v*_loc_ model are shown for each rightward hunting bout-type. Objects found in red bins increase probability of that bout-type being selected next. **F.** *pooled*-*i_n_* model summary. Activations of 5 bout-types are shown, given a range of *preceding interbout interval* values. Full bout-type probability distributions are shown for 3 specific values of *i_n_*(indicated with i, ii, and iii). For clarity, only probabilities of leftward bouts are shown, averaged across *fed* and *starved* fish. **G.** *pooled*-*t*_hunt_ summary. Similar to **F**, but for *hunt dwell-time*. **H.** *pooled*-*t*_explore_ summary. Similar to **F**, but for *explore dwell-time*. **I.** *split*-*t*_tank_ summary. Similar to **F**, but for *tank-time*. This feature is *split*, so *fed* and *starved* probabilities are shown separately. Note that *fed* and *starved* behavior is different at *t*_tank_ = 10 minutes, but converges by *t*_tank_ = 40 minutes.

### Simulating Behavioral Trajectories of *Fed* and *Starved* Fish

A strong test of a behavioral model is to evaluate its ability to generate realistic behavior in novel contexts. To that end, we alternate sampling from the *combo* bout and interbout models (here called a *combo renewal process*) to move a virtual fish through an artificial environment with abundant prey (**6A**, **Supplementary Videos 3-4**). We simulated behavioral trajectories of 50 *fed* and 50 *starved* fish and found that simulations capture multiple timescales of structure in larval zebrafish behavior. Prey object locations influence hunting bout selection in expected ways (**6B**), and *fed* fish select longer interbout intervals (**6C**), as expected. The effects of hunger can be seen by comparing bout transition probabilities of *fed* and *starved* fish in simulations (**6D**). For example, *fed* fish are more likely to transition to low-energy exploring bouts, while *starved* fish are more likely to transition to high-energy exploring bouts. *Starved* fish also adjust their behavior in several ways to increase food intake. They are more likely to transition to *J-turns* (especially *j2*) and to extend hunt sequences by linking *pursuits*^55^. *Starved* fish are also less likely to transition to *abort* and more likely to transition to *strike* than *fed* fish.

The *combo renewal process* simulations also capture longer timescale behavioral dependencies observed in our experiments. With real fish, we find that hunts ending in *strike* are much longer than those ending in *abort* (**6E**, left panel). This trend is absent in simulations generated from a *first-order Markov renewal process* (**6E**, middle panel) in which interbout intervals and bout-types depend on just *preceding bout-type* (*b_n−_*_1_). By contrast, the *combo renewal process* simulations do better to recover this higher-order hunting structure (**6E**, right panel). We also generate more realistic exploring behavior with the *combo renewal process*. Upon entering exploring mode, larvae are initially likely to exit, but get more locked into this mode with time (see **4H**). Accordingly, we see an overabundance of short and also long exploring sequences in *combo renewal process* simulations compared to simpler simulations (**6F-G**). Finally, the simulations capture slowly changing differences in bout-type selection probabilities across *fed* and *starved* fish as their hunger states equilibrate over 40 minutes (**6H**).

## Discussion

Behavior is the principal output of the nervous system and, like the nervous system, it is complex and high-dimensional^29, 61^. To make studying the brain more manageable, animal behavior is often constrained during neurobehavioral experiments. This reductionist approach has many benefits: it simplifies behavioral description, reduces experimental variability, and improves interpretability of neural data. However, an important frontier in neuroscience is to better understand how brains function in natural conditions^62^. The NIH BRAIN Initiative^63^ has identified study of the “Brain In Action”^64^ as a priority research area, stating that “[a] critical step ahead is to study more complex behavioral tasks and to use more sophisticated methods of quantifying behavioral, environmental, and internal state influences on individuals.”^65^ Continued development of such tools should allow for improved quantification of ethologically relevant behaviors occurring in naturalistic environments. Importantly, these tools should capture the dynamics of minimal behavioral elements, scale to big datasets, and be compatible with modern techniques to record neural population activity. Here, we describe such an approach for the study of larval zebrafish behavior.

In this study, we first acquired high resolution behavioral and environmental data of larval zebrafish exploring and hunting in an unrestrictive naturalistic environment. We then used dimensionality reduction and density-based clustering to categorize individual swim bouts into discrete types. Finally, we constructed a probabilistic generative model (specifically, a marked renewal process) that predicts future larval zebrafish behavior by combining information about hunger state, multiple timescales of behavioral history, and the positions and properties of potential prey in its local environment. This approach offers a means to summarize larval zebrafish behavioral patterns, to validate and compare models composed from separate predictive features, and to simulate realistic behavioral trajectories through virtual environments. By constructing a behavioral model of how state influences action, we can make predictions about what types of signals must be present in the neural system driving this behavior.

Our study generates testable hypotheses about how neural mechanisms might give rise to observed behavioral patterns on multiple timescales. On a short timescale, we see that larvae are likely to link consecutive exploring bouts through ipsilateral transitions, but also tend to begin hunts ipsilaterally. It has been shown that a reciprocally connected neural circuit in the anterior hindbrain (the ARTR) alternates between “leftward” and “rightward” states to mediate temporal correlations in turn direction during exploration^40^. However, how the ARTR or related circuits may bias reactions to prey stimuli is unknown. We predict the ARTR or related networks may asymmetrically modulate premotor systems (e.g. reticulospinal neurons^66^, hunting command neurons^67^, or premotor tectal assemblies^68^) such that, with the ARTR in a leftward state, leftward *J-turns* are generated preferentially to rightward *J-turns*. Alternatively, ARTR-state-dependant modulation could also occur further upstream in the sensorimotor hierarchy, for example, through asymmetric modulation of left- and right-hemisphere retinorecipient areas that process prey stimuli (e.g. optic tectum, AF7)^68, 69^. In this case, identical prey stimuli presented to the left or right eye may not be equally salient. Instead, output from the left and right retinas might be processed asymmetrically to promote ipsilateral transitions from exploring to hunting.

Following hunt initiation, several behavioral patterns interact to influence hunt outcome. As hunts extend, larvae select shorter interbout intervals, *pursuits* become finer, and *abort* probability decreases while *strike* probability increases. By contrast, as the time since the previous bout increases, *abort* probability increases, *pursuit* probability rises and falls, and *strike* probability decreases. We posit these time-sensitive hunting patterns depend on reciprocal connectivity between the nucleus isthmi (NI) and the tectum and pretectum^55^. It has been shown that the NI becomes active following hunt initiation and that NI ablation leads to specific deficits in hunt sequence maintenance, thereby increasing *abort* probability. Henriques et al.^55^ propose a model for NI-mediated feedback facilitation of (pre)tectal prey responses to facilitate extension of hunt sequences. This model can help explain why *abort* probability decreases and *strike* probability increases as hunts extend, though how bout-type selection during hunts depends so intricately on preceding interbout interval duration is not well understood. It is possible that tectal prey representations attenuate as the interbout interval extends beyond a few hundred milliseconds, potentially through phasic NI feedback that ramps and then decreases as each interbout extends. Alternatively, perhaps premotor populations^49,67,68^ involved in *pursuit* and *strike* generation become increasingly inhibited as interbouts get longer, increasing hunt termination probability. Since larvae tend to abort hunts through a contralateral bout transition, we suspect *abort* generation may frequently coincide with a change in ARTR state. Such a mechanism might facilitate switches in spatial attention from one hemifield to the other, thereby inhibiting return^70^ to a previously pursued object.

In our study, we observe many behavioral differences across *fed* and *starved* fish that equilibrate over tens of minutes. With respect to hunting, we see that *starved* fish are more likely to initiate and extend hunts. While engaged in a hunt sequence, *starved* fish also upregulate transitions to *strike* and downregulate transitions to *abort* relative to *fed* fish. It has been shown that food deprivation modulates larval zebrafish tectal processing of prey-like and predator-like visual stimuli such that food-deprived larvae are more likely to perceive and approach small moving visual objects^50^. Specifically, hunger induces recruitment of additional prey-responsive tectal neurons and neuroendocrine and serotonergic signaling mediates this effect^50^. We posit this mechanism may also increase tectal input to NI, thereby increasing NI-mediated feedback to facilitate hunt sequence extension and increased *strike* probability in *starved* fish. While not yet tested, direct hunger-state modulation of tectal-projecting NI neurons is also plausible. Other studies show that lateral hypothalamic neuron activity correlates positively with feeding rate^71^ and that lateral hypothalamic neurons respond to both sensory and consummatory food cues^72^. This brain region is likely critically involved in sustaining increased hunting over tens of minutes through modulating visual responses to prey and/or facilitating hunting (pre)motor circuits.

While the above mechanisms can explain why *fed* fish initiate fewer hunts in these experiments, we hypothesize satiety signals affect several other circuits to induce additional changes in exploring behavior. *Fed* fish select longer interbout intervals, lower-energy exploring bouts, and maintain wider eye divergence preceding all exploring bout-types. To coordinate these behavioral patterns, satiety cues may separately modulate midbrain nuclei involved in regulating bout timing^73^, nMLF neurons involved in regulating swim bout duration and tail-beat frequency^59^, and oculomotor centers involved in controlling eye position^74,75,76^. It is clear that feeding state coordinates a complex array of behavioral modifications, likely through modulation of many circuits distributed across the larval zebrafish brain.

There are many avenues to extend this work. In future studies of naturalistic larval zebrafish behavior, moving camera systems could employ faster camera frame rates, shorter exposure durations, and better tracking algorithms to improve raw behavioral data. These modifications will allow for higher resolution pose estimation (e.g. by including pectoral fin dynamics, pitch and roll estimates, and tail half-beat analysis^14^), facilitate more comprehensive bout-type classification, and yield significantly longer continuous behavioral sequences. Richer datasets will enable future models to extract nuanced environmental dependencies, like prolonged attention to single prey amongst many distractors during long hunt sequences (Bolton et al., manuscript in preparation). Such models might also combine behavioral history information with raw environmental video^77^ to sharpen behavioral predictions. Future models may simultaneously infer discrete behavioral states and their dynamics^8,15,78,79^, though the non-Markovian dependencies on past behavior present new challenges^80^. Likewise, there are many other internal state variables that could govern action selection, and future models could seek to infer these latent states^80^ rather than using proxy covariates like *tank-time*. Behavioral states and dynamics differ slightly from one individual larva to the next, and long-term behavioral recordings combined with hierarchical models^81, 82^ will allow us to study how these behavioral differences emerge and change throughout early development. In addition, the contribution of particular neural populations in generating naturalistic behavioral patterns may be probed by combining these behavioral models with experiments to activate, inhibit, or ablate specific neural cell-types in observed fish. Finally, our approach to modeling naturalistic larval zebrafish behavior may be combined with new technologies to record large neural populations in freely swimming fish^83,84,85,86,87^. This will present opportunities to construct joint models of neural activity and natural behavior, providing important tools to study the brain in action^88, 89^.

## Methods and Materials

### Animal Care

All fish were 7-8 days post-fertilization (dpf) Nacre^−/−^ zebrafish raised at 27C. Fish were given abundant live paramecia as food beginning at 5dpf. On test day, fish from the *fed* group remained in their Petri dish with abundant paramecia while fish from the *starved* group were placed in clean water for ~2-5 hours prior to testing. Testing was performed between 10 AM and 6 PM with 4-6 fish usually tested per day.

### BEAST Design

The gantry was acquired from CNC Router Parts (CRP4848 4ft × 4ft CNC Router Kit; www.cncrouterparts.com) and was modified to run upside-down on top of a support structure constructed from aluminum T-slotted framing available through 80/20 Inc (www.8020.net). Three electric brushless servo motors (CPM-MCPV-3432P-ELN ClearPath Integrated Servo Motors) and 3 Amp DC Power Supply (E3PS12-75) were acquired from Teknic (www.teknic.com). The camera (EoSens 3CL) was acquired from Mikrotron (www.mikrotron.de) with a frame-grabber from National Instruments (NI PCIe-1433; www.ni.com). The camera lens was acquired from Nikon (AF-S VR Micro-Nikkor 105mm f/2.8G IF-ED; www.nikonusa.com). A long-pass infrared filter was placed over the lens (62 mm Hoya R72; www.hoyafilter.com) to block light from the projector and collect transmitted light from an array of 16 IR-LED security dome lights (850 nm Wide Angle Dome Illuminators) positioned on the air table below the fish tank. The projector was acquired from Optoma (Optoma GT1080; www.optoma.com) and mounted on the side of the air table to project onto a diffusive screen (Rosco Cinegel 3026 Tough White Diffusion (216); www.stagelightingstore.com) embedded in the bottom of the plexiglass tank.

### Data Acquisition

The walls of the observation arena were assembled with light gray LEGO blocks to confine the fish to a water volume of 300 × 300 × 4 mm. Approximately 15 ml of water containing a high density of live paramecia were added near the center of the arena prior to testing each fish. This paramecia stock also contained some rotifers and algae particles. For testing, single fish were transferred to the arena where inwardly drifting concentric gratings were projected to bring the fish to the arena center. Zebrafish larvae tend to turn and swim in the direction of perceived whole-field motion, a reflexive behavior called the optomotor response, and we leverage this response to relocate the fish. Once the overhead camera detected the arrival of the fish, the first observational trial was initiated and the drifting gratings were replaced by a static color image of small pebbles, a natural image with reasonably high spatial contrast. Next, the camera moved automatically on the gantry to maintain position above the fish and capture video with a frame-rate of 60Hz and 2 millisecond exposure duration per frame. The fish was tracked for 3 minutes or until it reached the edge of the arena or tracking failed. The fish could evade the tracking camera with a high-speed escape maneuver, but these events were fairly rare. At the end of each trial, the camera returned to the arena center, video data was transferred from memory to hard disk, and concentric gratings were once again used to bring the fish back to the arena center to initiate another trial (up to 18 trials per fish). If the fish did not return to the center within 10-15 minutes, the experiment was terminated. The tracking algorithm was written to keep the darkest pixel in the image (usually contained within one of the eyes of the fish) within a small bounding box located at the image center. If the darkest pixel was located outside this bounding box, a command was sent to the motors to reposition the camera to the location of that darkest pixel. In this way, the camera moved smoothly from point to point to follow the fish, using the “Pulse Burst Positioning Mode” setting for the ClearPath motors. We run the ClearPath motors with Teknic’s jerk-limiting RAS technology engaged to generate smooth motion trajectories and minimize vibration during point to point movement. Experiments were run using PsychToolBox in Matlab.

### Image Registration and Fish Pose Estimation

In every image frame, connected component pixel regions corresponding to the left eye, right eye, and swim bladder were identified. The fish head center was defined as the average position of the centers of these 3 regions of interest. Heading direction is defined as the direction of the vector from the swim bladder center to the midpoint between the two eye centers. This information is used to translate and rotate each image for subsequent pose estimation and environment analysis. Only image frames in which all postural features could be extracted were included for further analysis. One common issue with pose estimation was caused by body roll of the fish, usually during an attempt to strike at a prey object, in which the fish would roll (rotate around its rostro-caudal axis) enough for one eye to occlude the other from the view of the overhead camera. Rather than estimate eye vergence angles in these situations, these image frames were excluded. Another common issue was caused by high-speed maneuvers by the fish during which a 2 millisecond exposure was insufficient to capture a suitably sharp image, thus causing either image registration or pose estimation algorithms to be compromised. We included only image frames in which all postural features could be accurately extracted for further analysis. Video segments containing problematic frames were split into separate video segments.

### Temporal Segmentation of Bout and Interbout Epochs

Bout and interbout epochs were identified by taking the absolute value of the frame to frame difference in heading angle and thresholding this time-series at 0.7 degrees. This binary signal was then dilated (radius = 2 elements) and eroded (radius = 1 element) with built-in Matlab functions (imdilate, imerode) to merge bout fragments and expand bout epochs to include one extra frame at the beginning and end of each bout. These operations set the minimal duration of both bout and interbout epochs to 3 frames (50 ms).

### Bout Summary Statistics

Δheading per bout: The change in heading angle per bout was calculated by averaging the heading direction of the fish over all frames in the preceding interbout epoch and subtracting this value from the average heading direction through all frames in the following interbout epoch. Positive values are assigned to leftward bouts.

|Δheading*|* per bout: The absolute value of Δheading per bout.

distance traveled per bout: The change in head position per bout was calculated by averaging the position of the fish head in the arena over all frames in the preceding interbout epoch (starting position) and also for the frames in the following interbout epoch (ending position). The distance between these two points is the distance traveled per bout.

|Δtail-shape| per bout: This non-negative 1-dimensional quantity summarizes how much the tail changes shape during the 10 frames used to represent each swim bout (as in **2B**). To compute this quantity, let the 10-frame tail angle measurements be placed in an array *T* with shape 20 × 10. First, the magnitude of frame to frame changes in each tail segment angle are computed to give a new array with shape 20 × 9. |Δtail-shape| is the sum of the absolute values of these 180 array elements. This value is divided by 180 to give units: radians per segment per frame. In Matlab syntax, this is computed as:

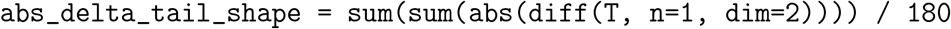

Several additional summary statistics are used to describe the fish eyes during each bout. These metrics are computed from the 10-frame representations of each bout and are described in S7.

### tSNE Input

As described in **2B**, each swim bout is represented as a 220-dimensional vector encoding the posture of the fish through 10 image frames beginning at bout initiation with 20 tail vectors and 2 eye vergence angles per frame (see also S1). All rightward bouts (Δheading *<* 0) were mirror reflected prior to running tSNE by swapping the left and right eye measurements and multiplying all tail angle measurements by Because tail measurements are tenfold more abundant than eye measurements and we sought to emphasize eye data in our clustering, we decreased the relative magnitude of the tail measurements by encoding tail measurements in radians and encoding eye measurements in degrees. In practice, we found that we could identify roughly the same bout-classes across a fairly wide range of scaling factors used to emphasize eye measurements relative to tail measurements. While it is common to preprocess data with PCA prior to running tSNE, we achieved qualitatively similar embedding results with and without this step, so we simply ran tSNE on the 220-D inputs.

Each of the 200,559 swim bouts are represented as 220-D vectors as described above and are embedded in a 2-D space with tSNE. We implement tSNE with Barnes-Hut approximations with CUDA to decrease tSNE runtime (https://github.com/georgedimitriadis/t_sne_bhcuda). Euclidean distance was used as a distance metric. Following embedding, 5 bout classes were identified with the routines described in S2. The 3 largest clusters were then further subdivided by kinematic variables, |Δheading*|* and |Δtail-shape*|*, as described in S6 and above.

### Identifying and Encoding Environmental Information

As described in **1K**, objects near the water surface preceding each swim bout are identified with image processing routines. Following image translation and rotation, a stack of the 6 images preceding bout initiation are cropped as in **1K**. High spatial contrast objects are identified in Matlab by filtering each image in this stack with a Laplacian of Gaussian 2-D filter (size = 13 × 13 pixels, sigma = 1.6). Contiguous 3-D object volumes in this image stack are smoothed with 3-D dilation and erosion. The average position of each object in the image frame preceding bout initiation was used to encode object location with the extrinsic feature *v*_loc_. The number of pixels assigned to each object in this final frame was used to encode object size with the extrinsic feature *v*_size_. The velocity of each object was computed by calculating the distance between the center of mass of each object in the 6^th^ image frame preceding bout initiation (t-minus 100 ms) and the 1^st^ frame preceding bout initiation (t-minus 17 ms) and dividing by the time elapsed. If an object was not properly segmented through all 6 frames, the velocity was calculated from fewer frames (with a minimum of 3 frames). The velocity vectors for each object were used to encode x-velocity and y-velocity in the extrinsic features *v*_x_ and *v*_y_. While the image resolution is sufficient to extract additional features such as the orientation, eccentricity, and detailed shape of each of object, we have not yet included this information in our predictive models.

### Intrinsic Features to Predict Interbout Intervals and Bout-types

#### preceding interbout interval

the duration of the preceding interbout interval in seconds. For models to predict interbout interval duration (*i_n_*), this is the duration of interbout *i_n−_*_1_. For models to predict bout-type (*b_n_*), this is the duration of interbout *i_n_*.

#### preceding bout-type

the category (i.e. bout-type) of the preceding swim bout. There are 18 bout-types, each of which is composed of left and right versions, giving 36 categories in total.

#### hunt dwell-time

the integer number of observed preceding consecutive hunting bouts (i.e. *J-turn*, *pursuit*, *abort*, *strike*). As hunt sequences extend, this value increases.

#### explore dwell-time

the integer number of observed preceding consecutive exploring bouts. For all predicted interbout intervals and bout-types, either *hunt dwell-time* or *explore dwell-time* will be zero, with the other being a positive integer. Only the contiguous bout sequence containing the predicted interbout interval (*i_n_*) or bout-type (*b_n_*) is used to define these feature values.

#### tank-time

the amount of time (in minutes) elapsed since the first trial was initiated for that fish.

### Extrinsic Features to Predict Bout-types

*v*_loc_: locations of potential prey in the local environment.

*v*_size_: sizes of potential prey in the local environment..

*v*_x_: relative x-velocities of potential prey in the local environment.

*v*_y_: relative y-velocities of potential prey in the local environment.

See S9-S10 for more information on intrinsic and extrinsic feature encoding.

### Simulations

For our *combo renewal process* simulations (Figure 6, **Supplementary Videos 3-4**), we simulated an environment with prey objects that move as biased random walking particles. Each particle has a constant size and speed. These particles influence bout-type selection as their locations, sizes, and relative velocities are encoded with the extrinsic features *v*_loc_, *v*_size_, *v*_x_, and *v*_y_. We simulated 50 *fed* and 50 *starved* fish for 40 minutes each (in 20 two-minute trials). Similar to our real experiments, prey are distributed with a centro-peripheral gradient, with the highest density of prey located near where trials begin. To simulate a behavioral sequence, an interbout interval duration is selected by randomly sampling from an interbout interval distribution generated from the *combo* interbout interval model (described in **5C**). Next, a bout-type is selected by randomly sampling from a bout-type probability distribution generated from the *combo* bout model (described in **5F**). Upon selection of a bout-type, we move the fish through its virtual world along the bout-type specific trajectories described in S14. Since the *combo* bout model depends on the environment in addition to behavioral history and hunger state, the virtual prey objects influence the fish’s behavioral trajectory. We call this combined bout and interbout model a *combo renewal process*. For comparison, we simulate 50 *fed* and 50 *starved* fish with a simpler model in which interbout intervals and bout-types depend only on *preceding bout-type* (*b_n−_*_1_). In these simulations, interbout interval durations are sampled from a probability distribution generated by the selected form of the *preceding bout-type* feature (*split*-*b_n−_*_1_), and bout-types are sampled from a bout-type probability distribution generated from the selected form of the *preceding bout-type* feature (*pooled*-*b_n−_*_1_). This model is referred to as a *first-order Markov renewal process*, and has no environmental dependencies. For both the *combo renewal process* simulations and the *first-order Markov renewal process* simulations, model weights are set at the maximum a posteriori (MAP) estimate of the GLM weights from the trained models.

**Figure 5:**
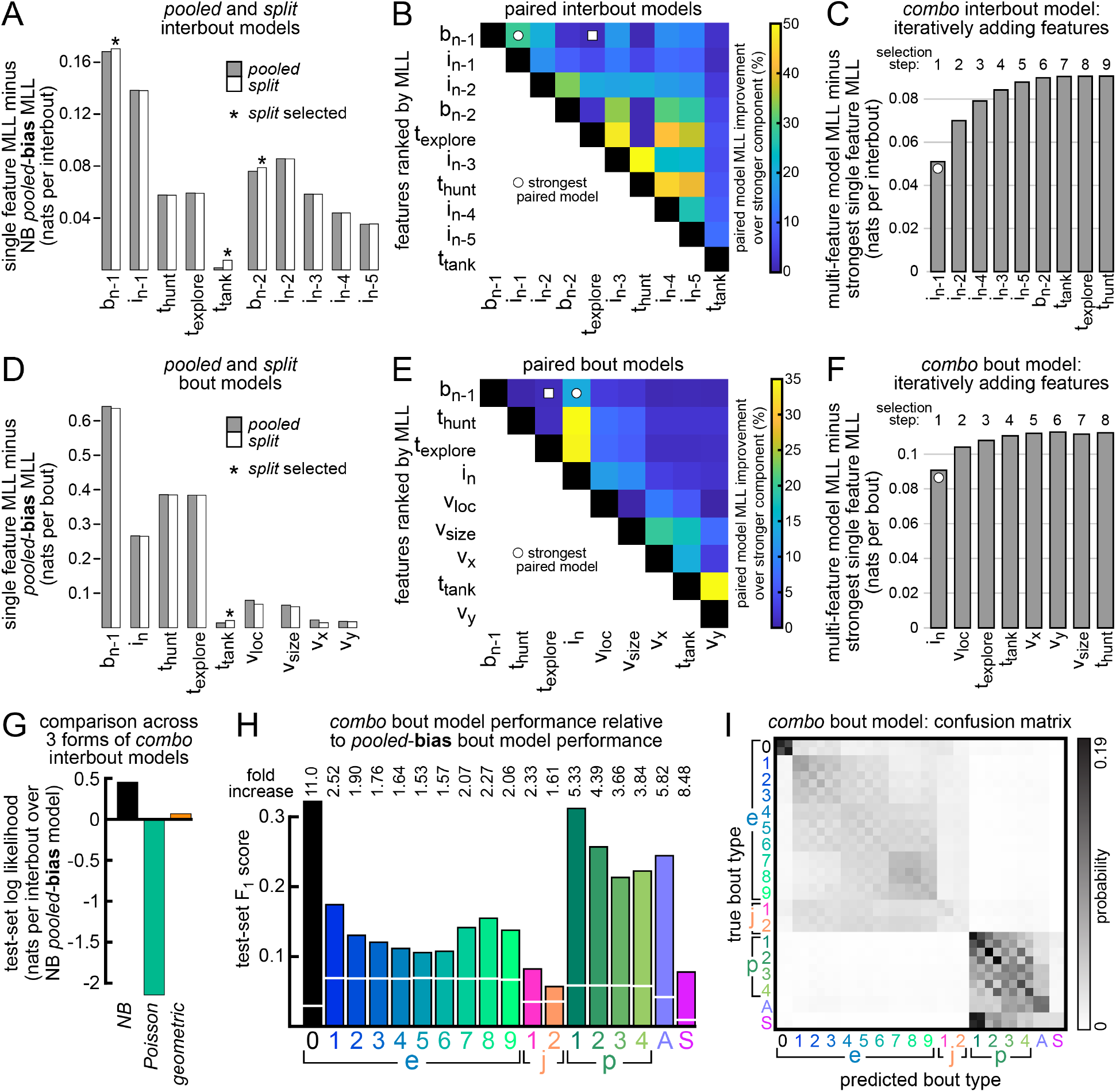
Comparing and combining interbout and bout models. **A.** *Pooled* and *split* forms of all *NB* interbout models introduced in Figure 3 are compared by computing the log of the marginal likelihood (MLL) of each model. MLL of the simplest interbout model (*pooled-bias*: 1 parameter for mean, 1 for variance) is subtracted from each model’s MLL as a baseline. For comparison, we include features similar to *preceding bout-type* and *preceding interbout interval*, but from further into behavioral history. *Pooled* forms are selected unless indicated with asterisk. **B.** All pairs of features (in selected form) from **A** are combined to make 45 paired interbout models. Paired models are evaluated by computing their MLL (above *pooled-bias*, as in **A**) and dividing this value by MLL of the stronger feature component (values from **A**). **C.** All features (in selected form) from **A** are combined to make a *combo* GLM for interbout intervals. We start with the strongest paired model and add the feature at each selection step which increases MLL most, until all features are added. **D-F.** The same as **A-C**, but for bout-type models. **G.** The performance of the *NB combo* interbout model is compared to similarly constructed *combo* interbout models that instead generate either *geometric* or *Poisson* distributions to predict *i_n_*. The *NB* distribution fits the data best. **H.** Performance of the *combo* bout model is compared to that of the simplest bout model (*pooled-bias*, capturing just overall abundance of each bout-type). The F_1_-statistic is computed for each bout-type on test-set data (*combo* model: bars; baseline: white lines). Fold increase is reported for each bout-type, which is just *combo* performance divided by baseline performance. Note that some bout-types are much harder to predict than others. **I.** *Combo* bout model confusion matrix for test-set data (left and right versions of bout-types are shown).

**Figure 6:**
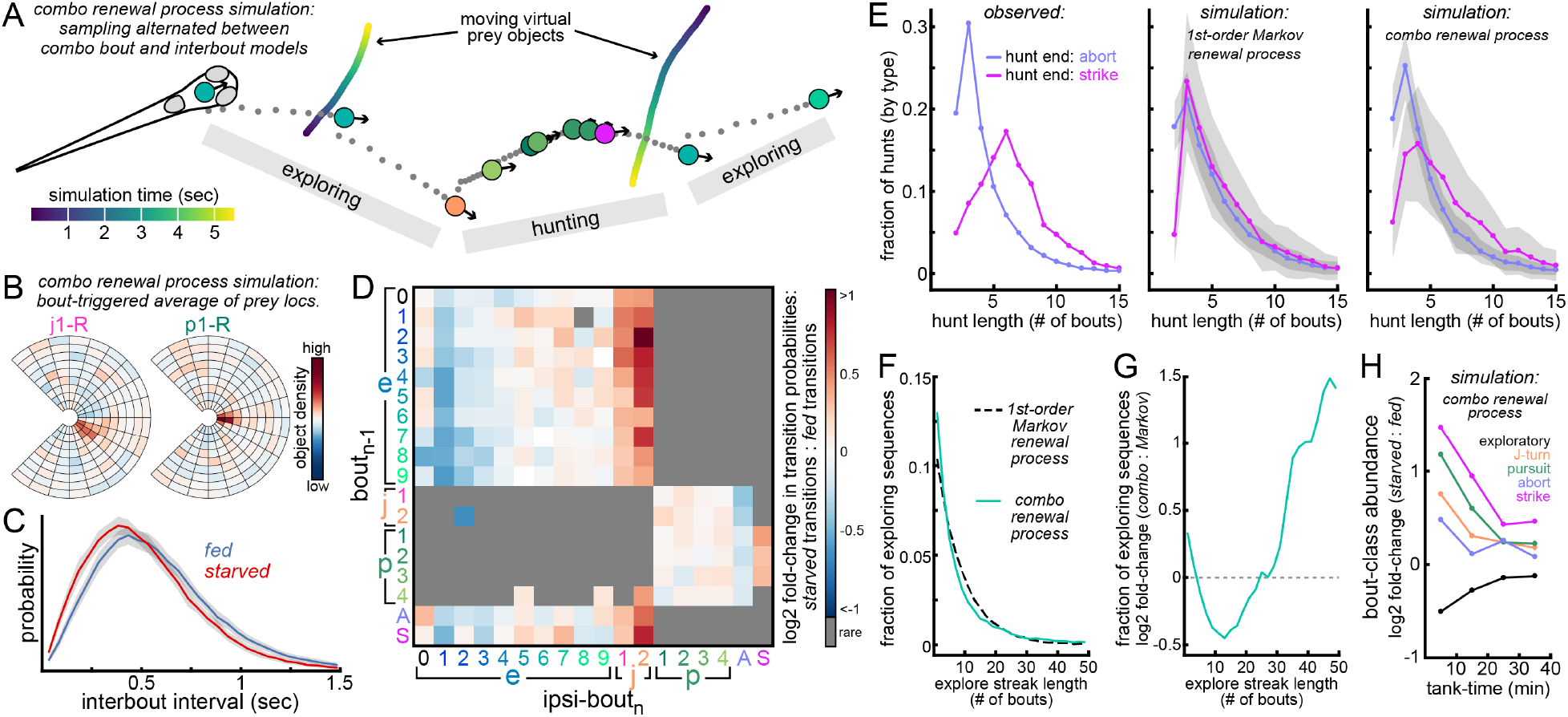
Sampling from *combo renewal process* to simulate behavior in a virtual environment. **A.** We simulate behavioral trajectories of 50 *fed* and 50 *starved* fish (40 minutes each) by alternately sampling from the *combo* interbout and *combo* bout models (we call this a *combo renewal process*). See **Supplementary Video 3-4**, S14, and Methods for details). Here, ~5 seconds of simulated behavior is shown in which the fish enters a hunt and *strikes* at a virtual prey object. Virtual prey objects influence bout-type selection and move through the environment as biased random-walking particles with fixed sizes and speeds. **B.** Bout-type triggered averages (for *j1-R* and *p1-R*) of the fish’s field of view preceding bout initiation indicate that prey objects influence hunting in expected ways. **C.** Histograms of interbout interval durations from simulated *fed* and *starved* fish (mean ± SE). **D.** Ipsilateral bout-transition probabilities observed for simulated *fed* and *starved* fish are compared. Red squares indicate increased probability for *starved* fish relative to *fed* fish. **E.** Hunt length distributions for hunts that end in either *strike* or *abort*. The left panel shows distributions for real fish observed in these experiments. The middle panel shows hunt length distributions (mean SE) from simulations in which bout-types and interbout intervals depend on only *preceding bout-type* (called a *1^st^-order Markov renewal process*). The right panel shows these distributions for simulations with the *combo renewal process* (as in **A**). **F.** Distribution of exploring streak length (in # of consecutive exploring bouts) for fish simulated from *1^st^-order Markov renewal process* and *combo renewal process*. **G.** Histograms from **F** are compared. *Combo renewal process* simulations produce more short (≤5 exploring bouts) and long (≥25 exploring bouts) streaks of exploring bouts. **H.** Bout-class abundances produced by simulated *fed* and *starved* fish (from *combo renewal process*) are compared in 10-minute bins (similar to **2D**).

## Data Availability

All data and modeling code will be made available upon final acceptance of the manuscript.

## Author Contributions

R.J. and F.E. conceived the project. T.P. and R.J. designed and constructed the BEAST. R.J. and C.W. conceived the basic experimental design. R.J. wrote the data acquisition code. R.J., C.W., and E.S. collected the data. R.J. processed and prepared all data for use in modeling studies. S.L. conceived and implemented all models with input from R.J. A.M. advised on neural network models. K.H. made the animations in **Supplementary Videos 1** and **3**. R.J. wrote the paper with input from all authors. S.L., C.W., K.H., and F.E. edited the paper. S.L. wrote the Technical Appendix.

## Acknowledgements

Research was funded by NIH grant U19NS104653 and Simons Foundation grant SCGB-325207. S.L. was supported by a Simons Collaboration on the Global Brain postdoctoral fellowship (SCGB-418011) and the Siebel Scholarship. We thank Ed Soucy, Joel Greenwood, and Adam Bercu of the Harvard Center for Brain Science Neuroengineering Core for technical support. We thank Andrew Bolton for helpful discussions on pose estimation and prey quantification, George Dimitriadis and Adam Kampff for assistance with GPU implementation of tSNE, and Matthew Johnson for helpful modeling discussions.

## Supplementary Figures

**Figure S1:**
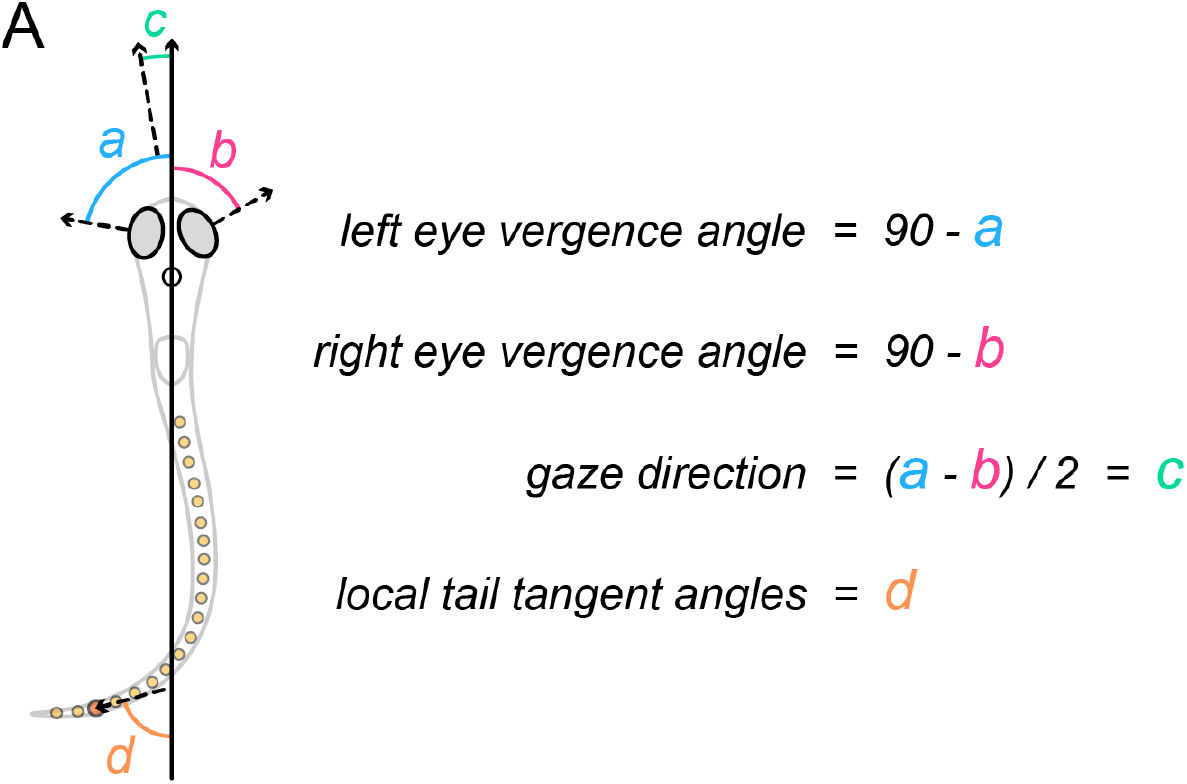
Encoding larval zebrafish posture in each image frame. **A.** In every image frame of our processed dataset, we estimate several postural features. Each image is first translated and rotated to align the fish head center (open black circle) and heading direction (solid black arrow) to a common reference. Left and right eye vergence angles are found by fitting an ellipse to each eye and comparing the direction of the minor axis of each ellipse to the fish heading (as shown). In this example, left eye vergence angle = 10° and right eye vergence angle = 30°. Eye vergence angles range from around −5° (highly diverged) to 45° (highly converged). Mean eye vergence angle is the average of the left and right eye vergence angles (20° in this example). Gaze direction is found by bisecting the angle formed between the minor axes of the eye ellipses, and quantifies the extent to which the fish is preferentially looking toward its left or right side. In this example, gaze direction = +10°. Gaze direction is positive for leftward gaze (as shown) and negative for rightward gaze. Tail shape is encoded in each image frame by placing 20 points along the tail and finding the local tail tangent direction at each point relative to heading, here shown for a single tail point. Tail tangent angles are positive when pointing toward the animal’s left side (shown at +75°) and are negative when pointing toward the animal’s right side.

**Figure S2:**
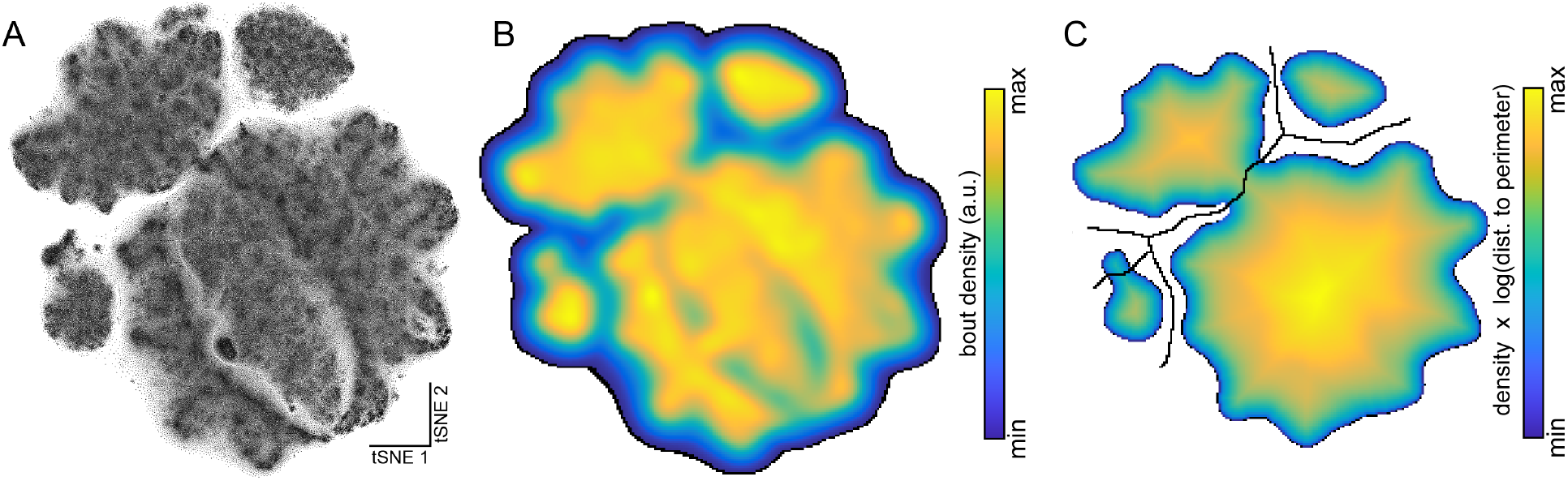
Identifying 5 bout-classes with density-based clustering of bouts in tSNE space. **A.** Each swim bout is represented as a 220-dimensional observation as described in **2B**. All rightward bouts (Δheading *<* 0) are mirror reflected to resemble leftward bouts. For reflection, left and right eye measurements are swapped and all tail vector measurements are multiplied by 1. All bouts are then embedded in 2-D space with tSNE (see Methods). **B.** The 2-D points in **A** are binned in a 200 × 200 pixel image and this image is blurred with a Gaussian kernel. **C.** The blurred image from **B** is thresholded to retain higher intensity pixels. Each pixel is then scaled by the log of the shortest distance from that pixel to the perimeter. A watershed algorithm is then implemented (black lines indicate segmentation boundaries) and bouts are assigned to 1 of 5 classes.

**Figure S3:**
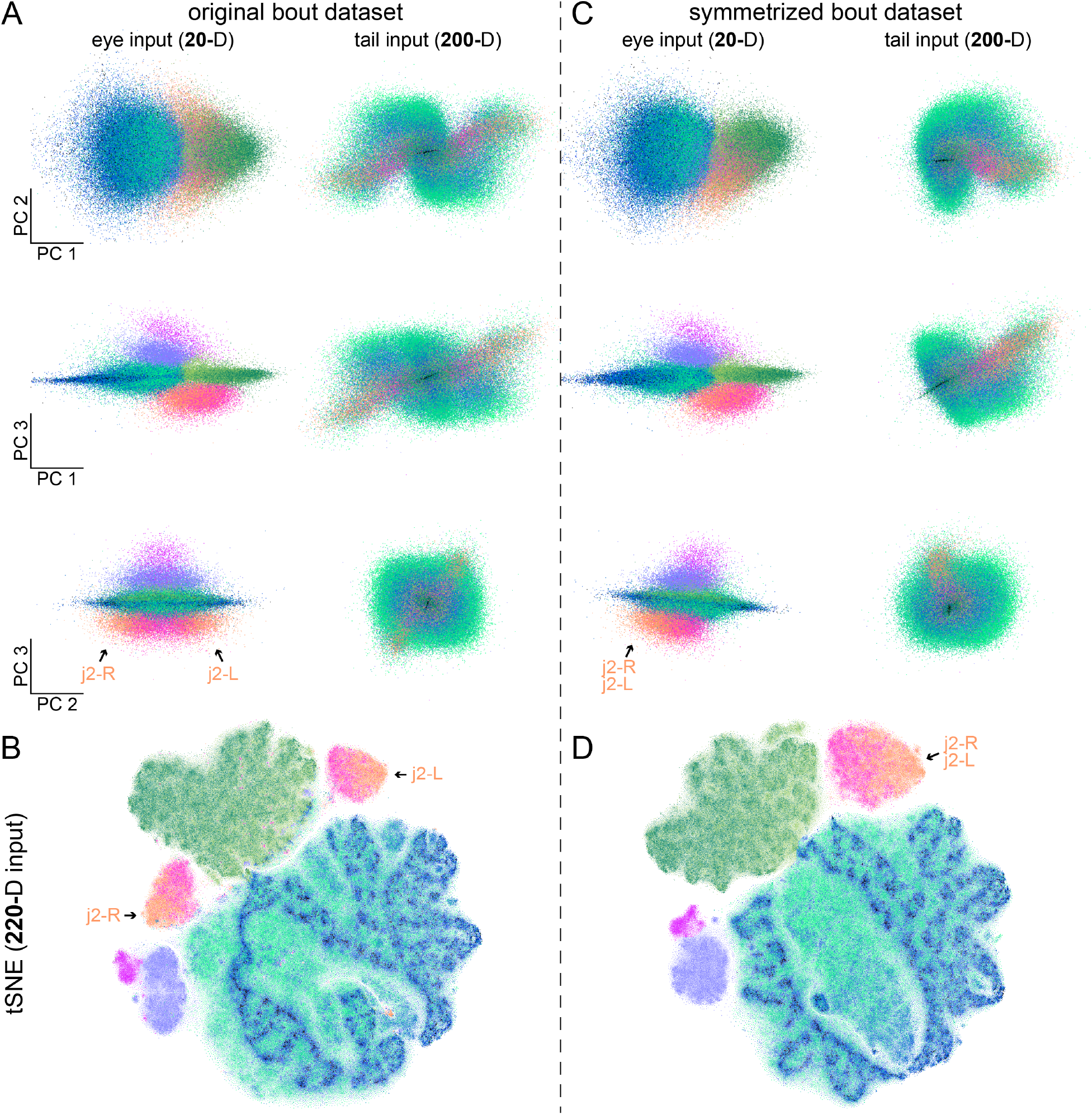
Exploiting bout symmetry to simplify dimensionality reduction. The bout dataset is highly symmetric, with approximately equal abundances of the left and right versions of each bout-type. Here we compare 2 types of dimensionality reduction (PCA and tSNE) on the original bout dataset (**A-B**) and the symmetrized bout dataset (**C-D**). **A.** Here we separately perform PCA on the original bout eye data (20-D representations, left column) and original bout tail data (200-D representations, right column). The first 3 principal components of the eye data and also the tail data are visualized, with each swim bout colored by its bout-type (as in **2E**). Bout-types are less intermixed in eye data PCA space than in tail data PCA space. The locations of left and right *j2* bouts are indicated. **B.** The original 220-D bout dataset is embedded in tSNE space. All tSNE parameters (including initial random seed) are identical to those used in the main text (**2C**). Note the separate clouds for left and right *J-turns*, which would complicate subsequent density-based clustering. **C-D**. Similar to **A-B** but for the symmetrized bout dataset. Note that all *J-turns* are now merged in a single cloud in **D**.

**Figure S4:**
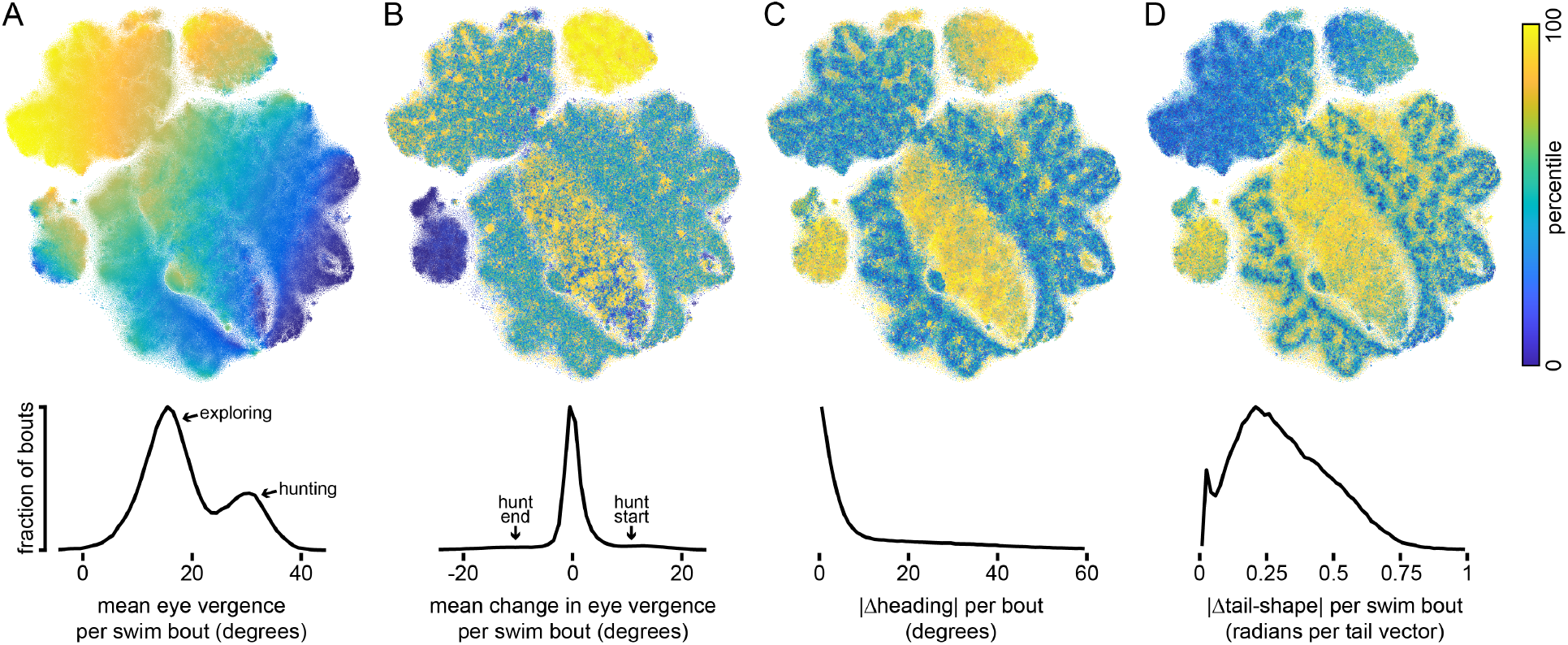
Visualizing bout dataset with kinematic parameters. The tSNE map from **2C** is reproduced in **A-D**. Each point is a swim bout. The color of each point is used to indicate its percentile for a given kinematic parameter. Histograms for these kinematic parameters for all swim bouts are shown in the bottom row. **A.** Mean eye vergence per swim bout, calculated from the 20-D eye representations of each swim bout. **B.** Mean change in eye vergence per swim bout, calculated from the 20-D eye representations. The mean eye vergence in frame 1 is subtracted from the mean eye vergence in frame 10. Note that larvae converge their eyes during *J-turns* (yellow cloud) and diverge their eyes during *aborts* and *strikes* (blue cloud). The 20-D eye observations largely influence the global structure of the tSNE map. **C.** Magnitude of change in heading angle per swim bout. This is calculated by comparing the average heading direction in the interbout interval preceding and following each swim bout. **D.** Magnitude of change in tail-shape (Δtail-shape) per bout is calculated from the 200-D tail observations for each swim bout (see Methods).

**Figure S5:**
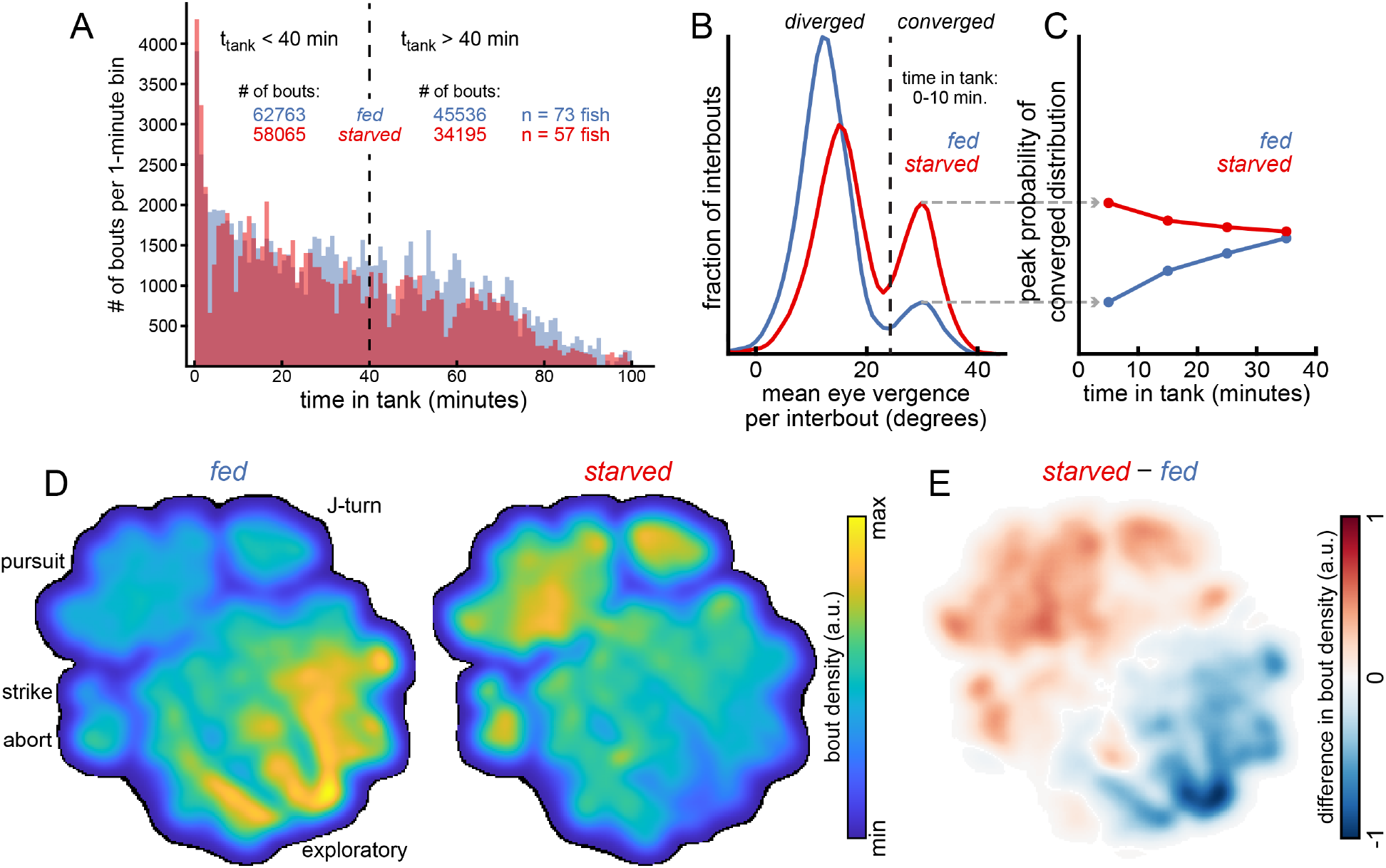
Hunger state modulates eye position and bout-type selection. **A.** Histogram of time elapsed since initiation of first observational trial (*tank-time*, *t*_tank_) for all observed bouts across *fed* and *starved* fish groups. 60% of all swim bouts occurred in the first 40 minutes (*t*_tank_ *<* 40 min) of testing and these bouts are used to compare *fed* and *starved* fish groups in **2D** and **2G-H**. The whole dataset is used for all modeling efforts and so include data with *t*_tank_ *>* 40 minutes. **B.** Reproduction of **2A** from main text. **C.** Histograms of mean eye vergence per interbout are calculated as in **B**, but for each 10-minute time bin to include all bouts with *t*_tank_ *<* 40 minutes. Each of these histograms is bimodal, and we report the height of the peak associated with eye convergence. The heights of these peaks converge as *tank-time* increases, demonstrating how *fed* and *starved* fish groups adjust their behavior over time. **D.** We randomly select 50,000 swim bouts (with *t*_tank_ *<* 40 min) from each fish group, plot these points in tSNE space, and make separate density maps for *fed* (left panel) and *starved* (right panel) groups as in S2B. **E.** The image in the left panel of **D** is subtracted from the image in the right panel of **D**. Red regions indicate bouts that are upregulated by *starved* fish.

**Figure S6:**
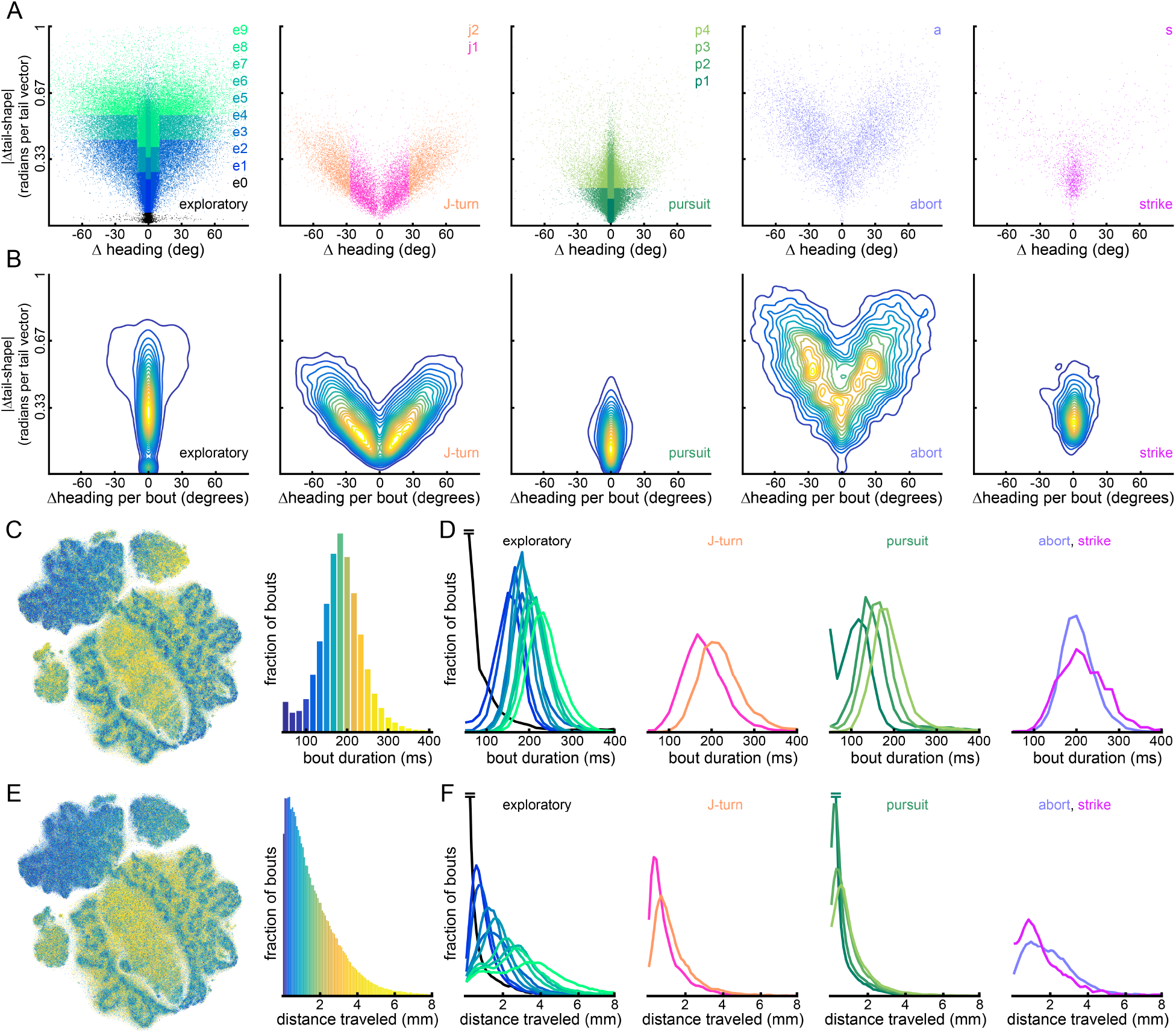
Subdividing bout-classes by kinematic parameters to get bout-types. **A.** For each bout-class, we plot |Δtail-shape| versus |Δheading| for all swim bouts in that class (see Methods). All bout-class distributions are highly symmetric. As described in the main text, the *exploratory*, *J-turn*, and *pursuit* bout-classes are subdivided. The *J-turn* class is split in half by |Δheading| to give *j1* (|Δheading| *<* 26.6°) and *j2* (|Δheading| *>* 26.6°). The *pursuit* class is larger and subdivided into 4 bout-types by first splitting all *pursuits* in half by |Δheading|, and then splitting each remaining group in half by |Δtail-shape|. Bout-types *p1* and *p3*: |Δheading| *<* 3.1°. Bout-types *p2* and *p4*: |Δheading| *>* 3.1°. Bout-types *p1* and *p2*: low-|Δtail-shape|. Bout types *p3* and *p4*: high-|Δtail-shape|. The *exploratory* class is the largest and the *e0* bouts (mostly orofacial and pectoral fin movements, plus some noise) are isolated first by thresholding at a local minimum in the bimodal distribution for |Δtail-shape| for this bout-class. Bout-types *e1*, *e4*, and *e7*: |Δheading| *<* 2.4°. Bout-types *e2*, *e5*, and *e8*: |Δheading| *>* 2.4° & *<* 10.3°. Bout-types *e3*, *e6*, and *e9*: |Δheading| *>* 10.3°. Bout-types *e1-3*: low-|Δtail-shape|. Bout-types *e4-6*: medium-|Δtail-shape|. Bout-types *e7-9*: high-|Δtail-shape|. **B.** Density contour plots derived from scatter plots in **A** with isolines delineating 5% quantiles. **C.** Each bout in tSNE map is colored by its percentile for bout duration (left panel). The histogram for bout duration is shown with corresponding color-code. **D.** Bout duration histograms are presented for each bout-type. All subplots share a vertical axis. Horizontal lines (on *e0* histogram) indicate data is truncated for display. **E-F.** Similar to **C-D** but for distance traveled per bout (see Methods).

**Figure S7:**
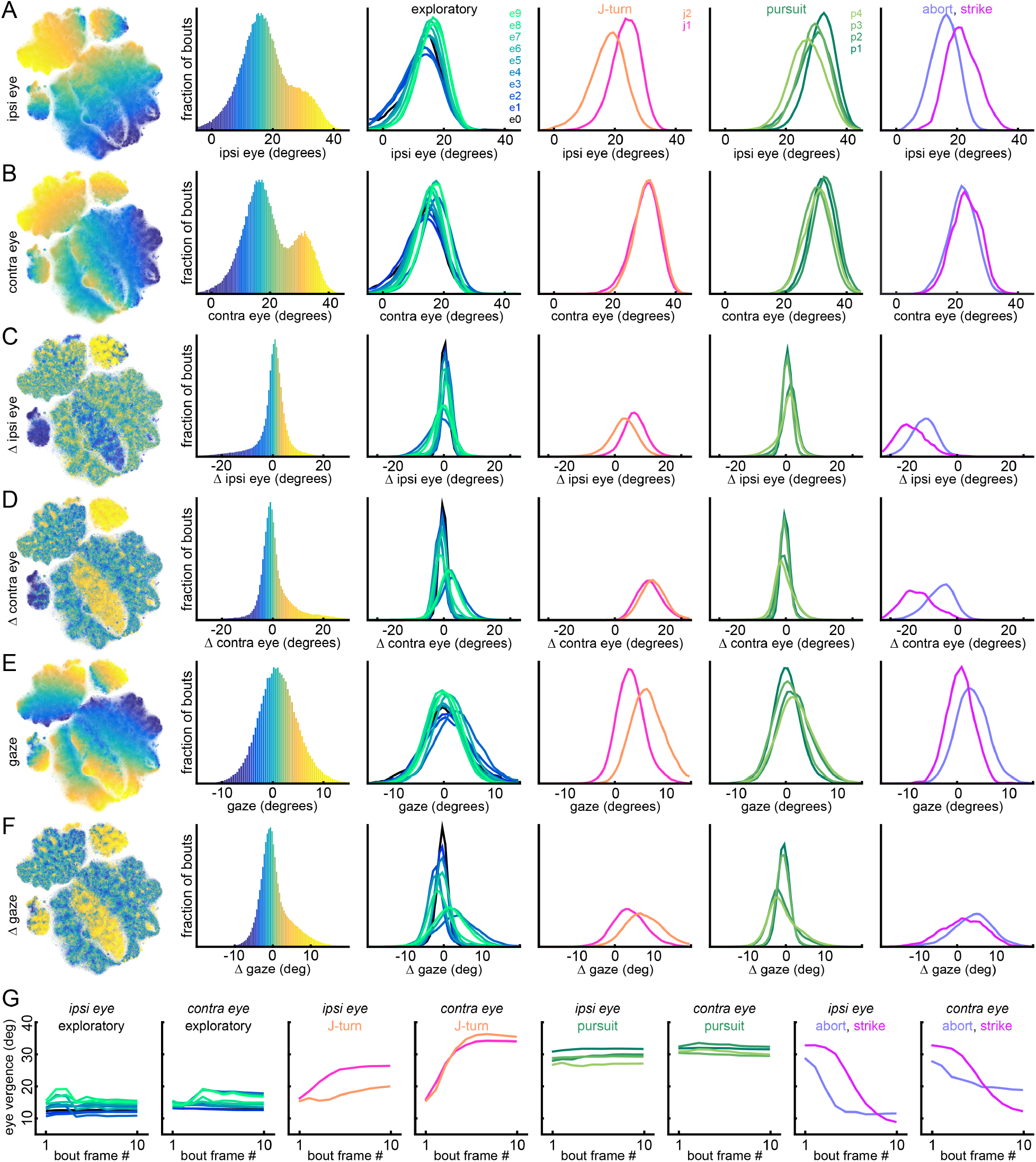
Eye movement patterns across bout-types. Each row in **A-F** is similar to S6C-D. Each swim bout is colored by its percentile for a given parameter summarizing the 20-D eye input data (panel 1 of each row). A histogram summarizes the distribution of this parameter over all bouts (panel 2 of each row) and provides the color reference for the corresponding tSNE map. Histograms are also shown for each bout-type (panels 3-6 of each row). **A.** Mean vergence angle of the ipsilateral eye across 10 image frames beginning at bout initiation. For a leftward swim bout, the ipsilateral eye is the left eye. **B.** Mean vergence angle of the contralateral eye across 10 image frames. **C.** Change in ipsilateral eye vergence angle. This is computed as the vergence angle in frame 10 minus the vergence angle in frame 1. **D.** Change in contralateral eye vergence angle. **E.** Mean gaze direction across 10 image frames. In each frame, gaze direction = (ipsilateral eye vergence angle contralateral eye vergence angle) */* 2 (see S1). Note that most *J-turns* have a positive gaze direction, as larvae look in the direction to which they will turn. **F.** Change in gaze direction. This is computed as gaze direction in frame 10 minus gaze direction in frame 1. Note that most *J-turns* occur with positive gaze change as the contralateral eye converges more than the ipsilateral eye, especially for *j2*. High-|Δheading| exploring bouts (*e3*, *e6*, *e9*) and *aborts* also tend to occur with large positive gaze change. **G.** Bout-type averages of ipsilateral and contralateral eye vergence angles across 10 image frames, beginning at bout initiation. Note the different eye movement patterns during *abort* and *strike*. During *strike*, larvae maintain eye convergence for longer, allowing them to maintain binocular view of a prey target as they attempt to capture it. Eye movement patterns during *strike* are also more symmetric than during *abort*.

**Figure S8:**
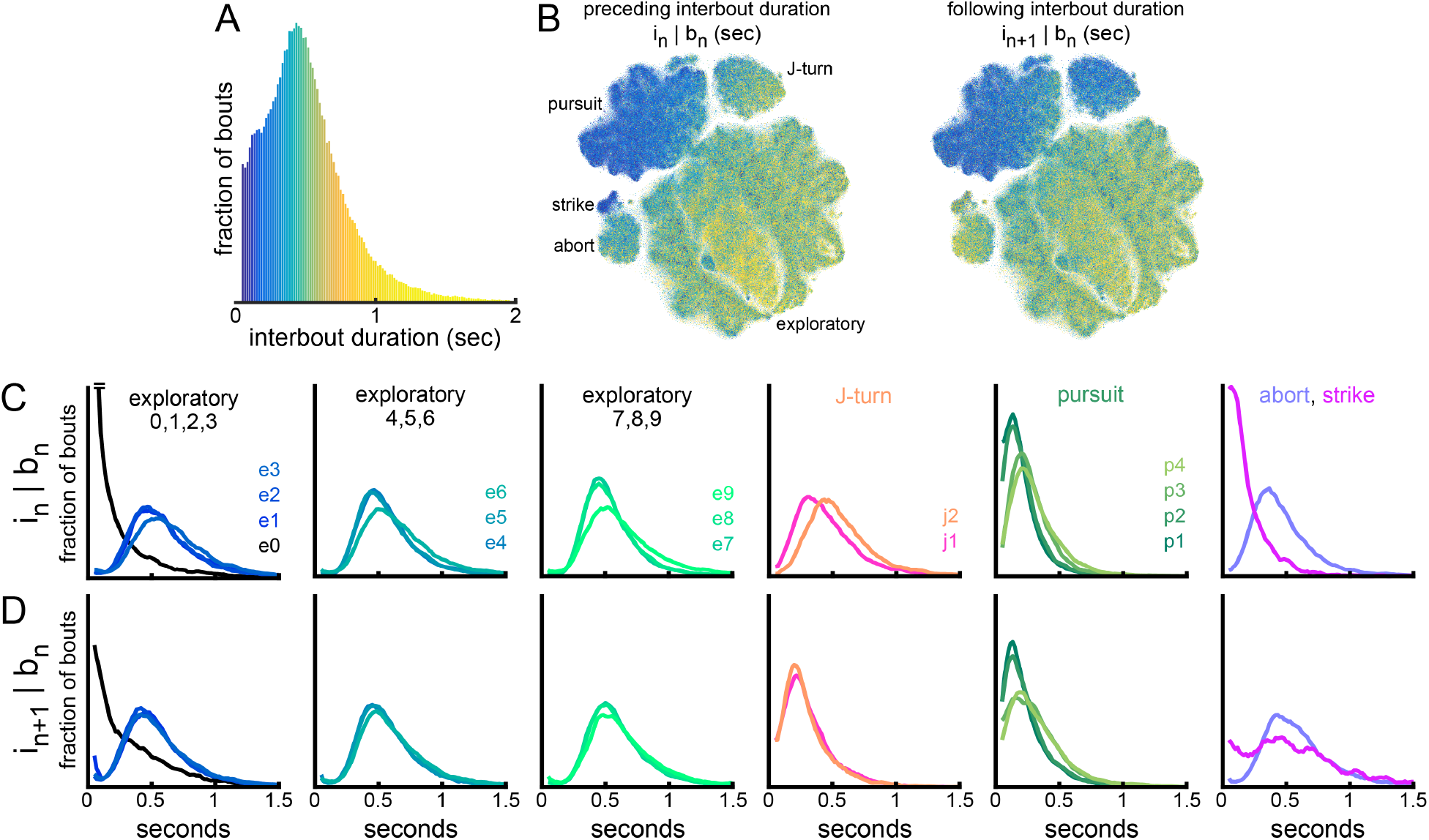
Duration of interbout intervals preceding and following each bout-type. **A.** Histogram of interbout interval duration for whole dataset. **B.** Each bout in tSNE space is color coded by the duration of its preceding (left panel) and following (right panel) interbout interval. Histogram in **A** provides the colormap. **C-D.** Histograms for duration of interbout intervals preceding and following each bout-type. All panels share a vertical axis.

**Figure S9:**
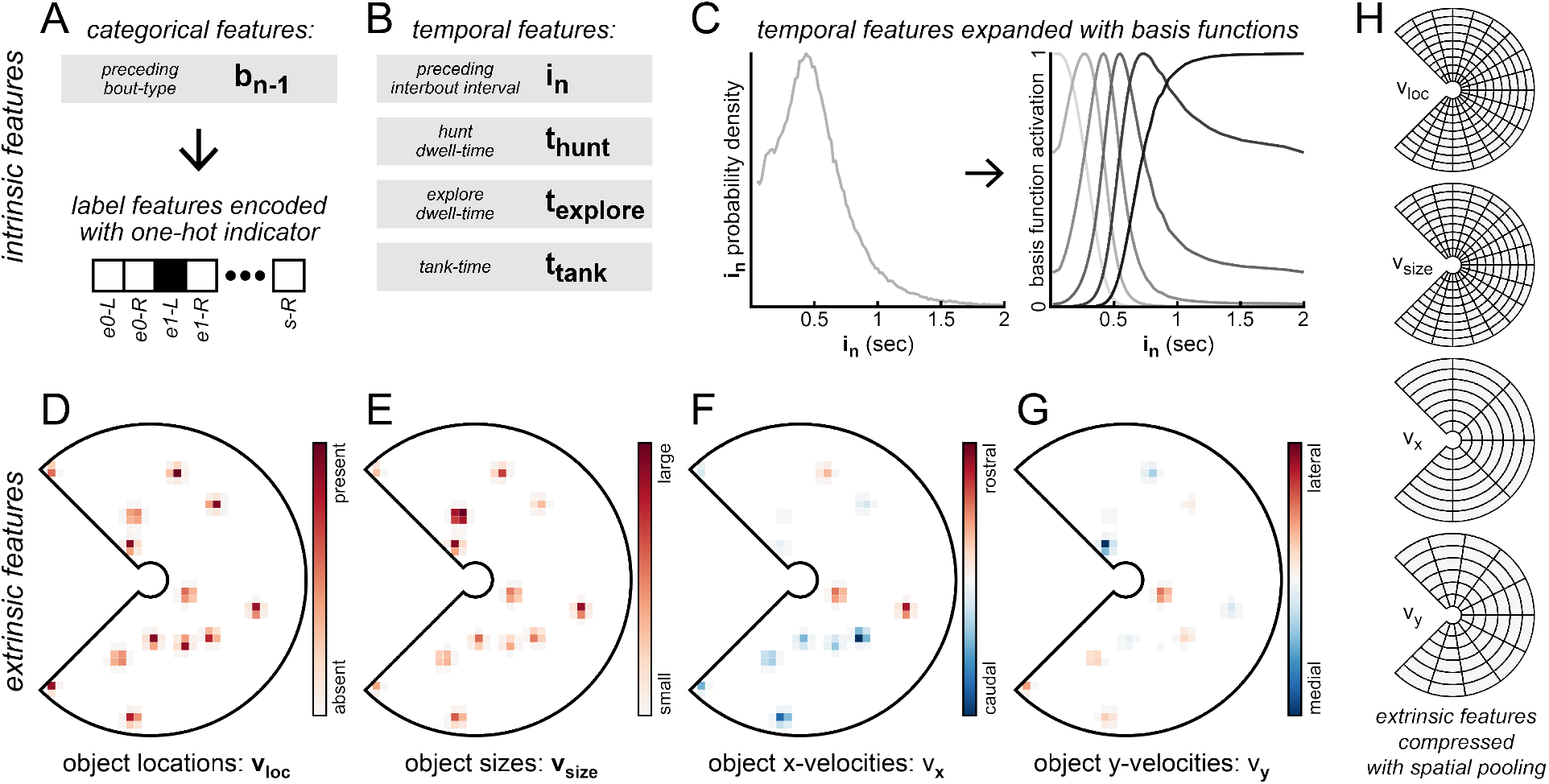
Encoding intrinsic and extrinsic predictive features with basis functions. Intrinsic features encode information about internal state or behavioral history and come in two categories: label features and temporal features. Intrinsic features are used to predict interbout intervals (*i_n_*) and bout-types (*b_n_*). **A.** Label features include *preceding bout-types* (*b_n_*_−1_ or *b_n_*_−2_) and are encoded with a one-hot indicator. **B.** Temporal features are 1-dimensional and are expanded to multiple dimensions with radial basis functions, allowing GLMs to learn non-linear mappings from a 1-D input to a probability distribution over *i_n_* or *b_n_*. Temporal features include *preceding interbout intervals* (*i_n_*, *i_n_*_−1_, *i_n_*_−2_…), *hunt dwell-time* (*t*_hunt_), *explore dwell-time* (*t*_explore_), and *tank-time* (*t*_tank_). **C.** The distribution of each 1-D temporal feature defines the shape of radial basis functions used to encode that feature (see Technical Appendix). For example, a given value for *preceding interbout interval* can be encoded as a 6-dimensional input by evaluating each of 6 basis functions at that input value. We select the number of basis functions used to encode each temporal feature with an empirical Bayes hyper-parameter grid search (S10). **D-G**. Extrinsic features encode information about objects in the local environment just prior to bout initiation and are used to predict that upcoming bout-type (*b_n_*). Here we show an example for encoding the locations, sizes, and relative velocities of each surface object found in **1K** to represent environmental state preceding initiation of the swim bout shown in **2B** (a *p4-R* bout). **D.** The center of mass of each object in the image frame preceding bout initiation (t-minus 17 ms) is used to encode the 868-dimensional *v*_loc_ feature by representing each object with a circularly symmetric 2-D gaussian (*σ* = 105 µm). The value of each of these 868 pixels is the value of the gaussian evaluated at the distance from the pixel to the object center, summed over evaluations for all objects. If no objects are present, all *v*_loc_ pixel values are zero. The *v*_loc_ feature is adjusted to generate the other extrinsic feature inputs. **E.** To encode object sizes with *v*_size_, the magnitude of the gaussian representing each object is scaled proportionally to its pixel area. Note how the large dust particle (box 1 in **1K**) is represented with high-intensity pixels while smaller objects have lower-intensity representations. **F.** The x-axis velocity component for each object is encoded with *v*_x_ by multiplying each gaussian by a factor proportional to that object’s relative x-axis velocity. Objects moving in the rostral direction (to the right) are assigned positive values while objects moving in the caudal direction (to the left) are assigned negative values. Note that the motionless dust particle is assigned near-zero value pixels while several motile prey objects are assigned non-zero value pixels. **G.** The y-axis component of relative velocity of each object is encoded with *v*_y_. This component is encoded symmetrically such that objects moving laterally (away from the x-axis) have positive values and objects moving medially have negative values. **H.** Extrinsic features are high-dimensional and we compress them to lower-dimensional representations by summing over pixel values within spatial bins. We select the number of radial and angular bins for each feature with an empirical Bayes hyper-parameter grid search (S10). The bin structure selected for each feature is shown.

**Figure S10:**
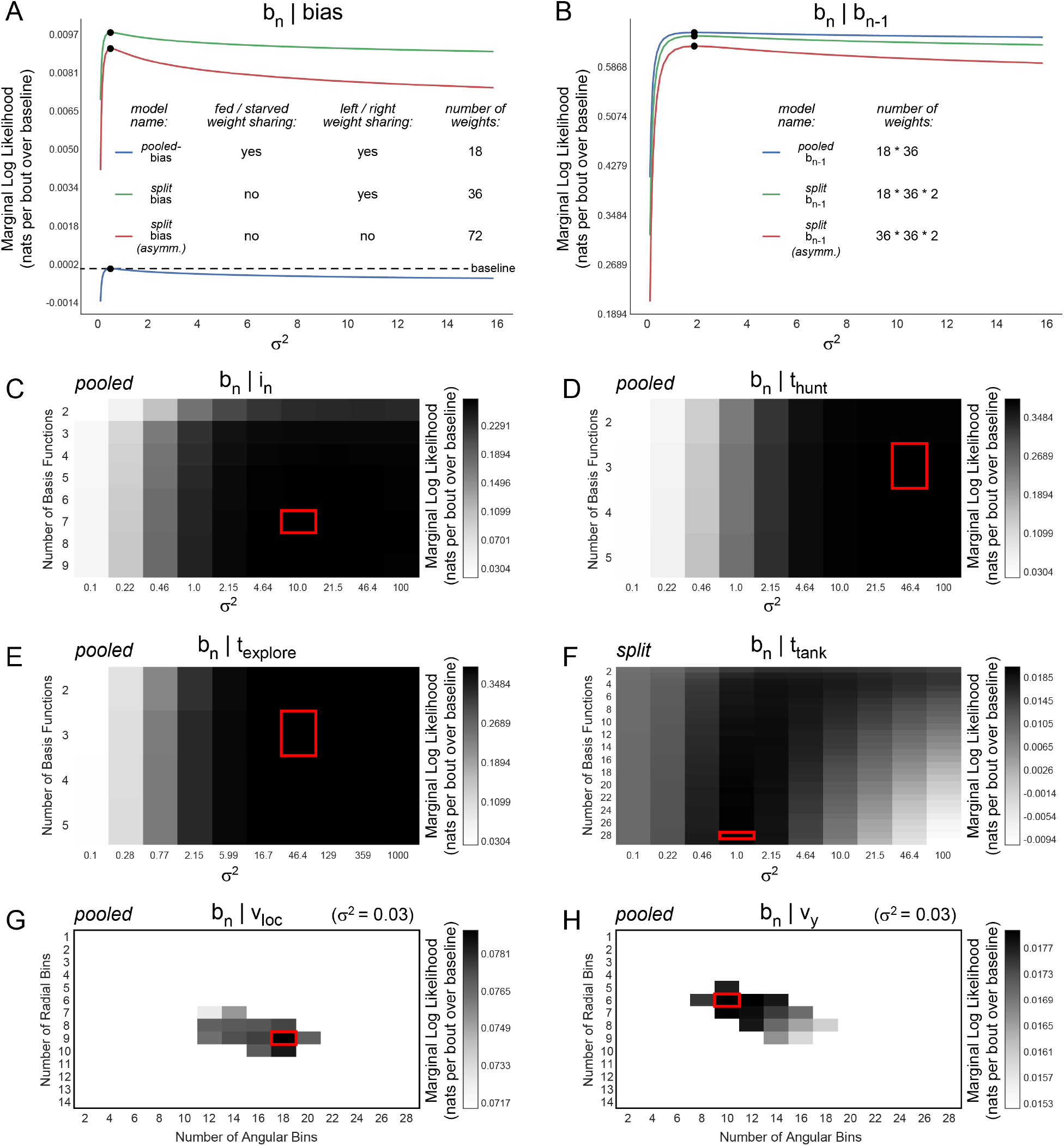
Empirical Bayes hyper-parameter selection. Here we describe our approach to perform empirical Bayes hyper-parameter selection for features used to predict bout-type *b_n_*. We take a similar approach to tune our interbout interval models (selection process not shown) **A.** The marginal log likelihood (MLL) of 3 forms of the *bias* model to predict bout-type are compared. The form that performs best (green line) is symmetric (one weight is shared for left and right versions of each bout-type) and *split* (separate weights for *fed* and *starved* fish groups). For each form, we sweep over values for prior variance (*σ*^2^) on the weights (selection indicated with black circles). The symmetric *pooled-bias* model (blue line) is used as a comparative baseline (dashed line) for all bout models, and its MLL (in nats per bout) is subtracted from MLL (in nats per bout) of other bout models to report their quality (in **B-H**, and also in main text). All bout models that take a predictive feature as input (except *preceding bout-type*) also include a set of asymmetric *split-bias* weights (red line) to account for bout-type abundance. **B.** Prior weight variance is selected for each of 3 forms of the *preceding bout-type* model to predict *b_n_*. The selected form is symmetric and *pooled* (*pooled*-*b_n_*_−1_: blue line), though it is only slightly favored over the symmetric and *split* model (*split*-*b_n_*_1_: green line), which has twice as many weights. The asymmetric form (*split*-*b_n_*_−1_: red line) is shown for comparison only, and is not considered independently in the main text. **C-F.** For each intrinsic temporal feature, we need to select both the number of basis functions used to encode the feature input (y-axis) and the prior variance (*σ*^2^) on the weights (x-axis). We perform a grid search and select the combination of parameters which gives the highest MLL (indicated with red box). For each feature, we perform a grid search for the *pooled* and the *split* form. We select the *pooled* form for all features to predict *b_n_*, except for *t*_tank_. **G-H.** For extrinsic features, we select 3 hyper-parameters (searches for *v*_size_ and *v*_x_ not shown). We select the number of radial (y-axis) and angular bins (x-axis) used to encode environmental objects (see S9H), and also the prior variance (*σ*^2^) on the weights. Since this search space is larger, we perform a greedy search. To do this, we compute MLL for a single combination of parameters. We then compute MLL for the 6 “neighboring” models (in 3-D parameter space), and pick the strongest parameter combination. We repeat this process until MLL is maximized (indicated with red box).

**Figure S11:**
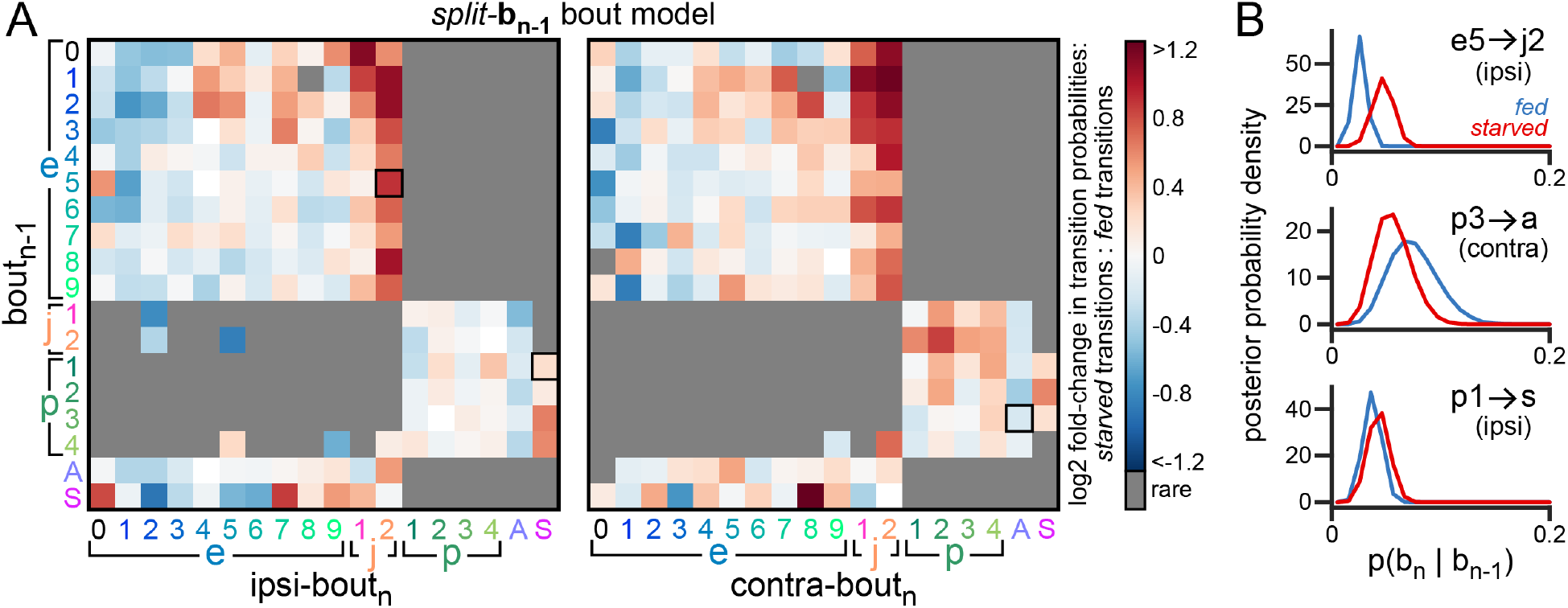
Comparing *fed* and *starved* bout transition probabilities with *split*-*b_n_*_−1_ model. **A.** Output probabilities of the *split*-*b_n_*_−1_ bout model. For each transition, transition probability for *starved* fish is divided by transition probability for *fed* fish, and the log_2_ of this quantity is reported. For clarity, rare transitions (<1% emission probability) are not shown. Blue squares show transitions with increased probability for *fed* fish, and red squares show transitions with increased probability for *starved* fish. While the *pooled*-*b_n_*_−1_ bout model was slightly preferred to this *split*-*b_n_*_−1_ form, this *split* form is useful to compare the behavior of *fed* and *starved* fish. The *pooled* and *split* form of this *preceding bout-type* model have nearly equal values for MLL. We expect potential gains in predictive performance with the *split* form are offset by increased complexity (it has twice as many free parameters). **B.** The posterior probability density for 3 selected bout transitions (black boxes in **A**) for *fed* and *starved* fish. These distributions were computed by sampling 10,000 sets of *split*-*b_n_*_−1_ weights and calculating output transition probabilities under each model instance. Note that *starved* fish are more likely to transition ipsilaterally from *e5* to *h2*, are less likely to transition contralaterally from *p3* to *abort*, and are more likely to transition ipsilaterally from *p1* to *strike*. While many bout transitions have significantly different probabilities in *fed* and *starved* fish, many also have similar probability or are rarely observed. This highlights some of the challenges of model selection because while the *pooled*-*b_n_*_−1_ bout model has fewer weights and slightly higher MLL, the *split*-*b_n_*_−1_ bout model captures many differences across *fed* and *starved* groups. Fortunately, our *combo renewal process* simulations (Figure 6) recover most of the differences in bout transition probability across *fed* and *starved* fish due to inclusion of other features with *split* architecture.

**Figure S12:**
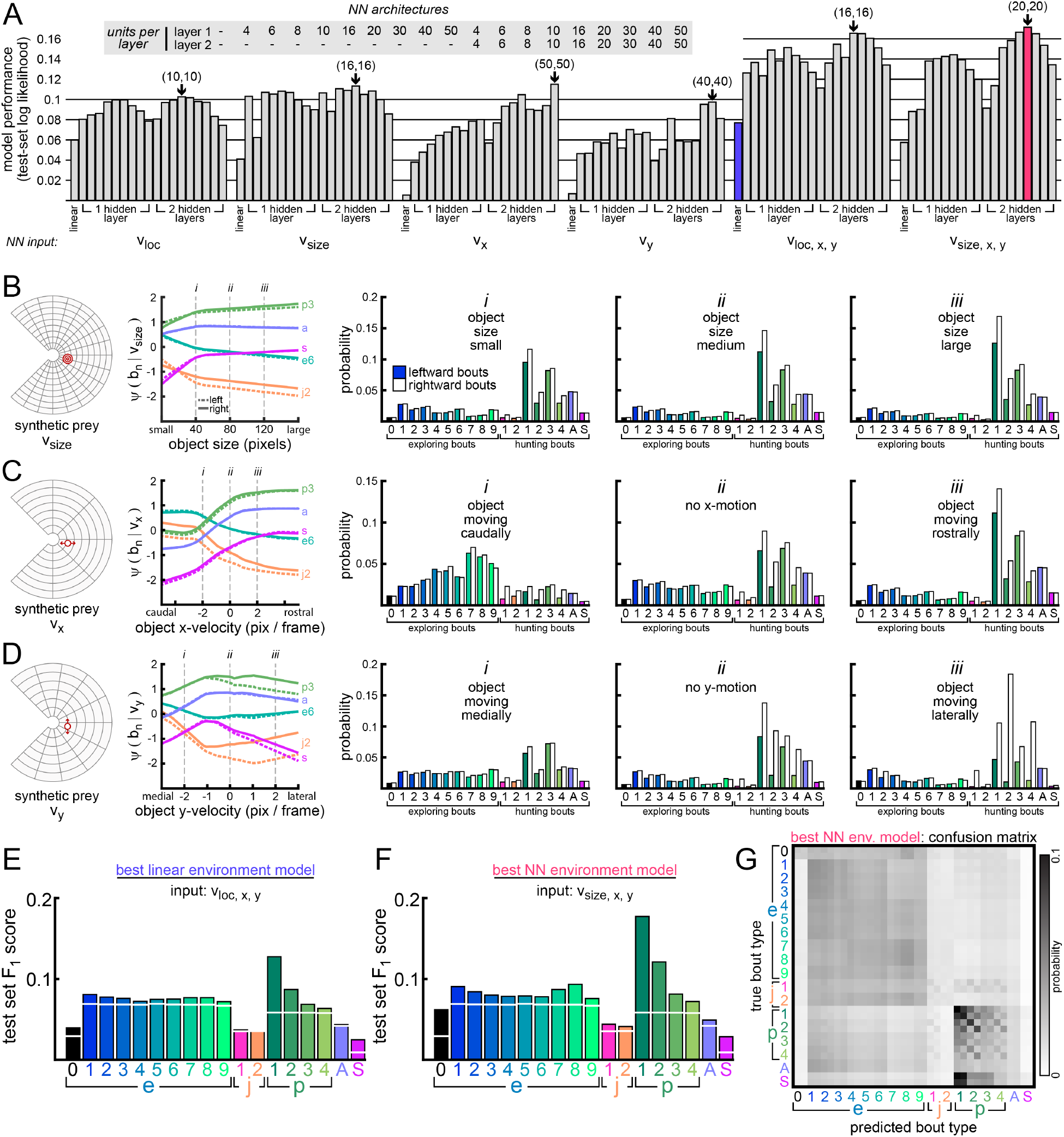
Feed-forward neural networks improve bout-type prediction from extrinsic features. **A.** Here we improve bout-type prediction from environmental information encoded in our extrinsic features (*v*_loc_, *v*_size_, *v*_x_, *v*_y_) by using feed-forward neural networks (NN) that take these features as input. We consider NNs with 1 or 2 hidden layers, with 4, 6, 8, 10, 16, 20, 30, 40, or 50 units per hidden layer. Layers are densely connected. We fit all NN architectures for each individual extrinsic feature input (blocks 1-4), but also consider combinations of extrinsic feature inputs (blocks 5-6). For comparison, we include GLMs described in **5D** of the main text (indicated with “linear” in blocks 1-4). The best model in each block is indicated (black arrow). Models are compared by the average log likelihood assigned to target bout-types in a held-out test-set. The strongest linear model (purple bar) and NN model (red bar) are indicated. The strongest NN model has 2 hidden layers, 20 units per layer, and takes *v*_size_, *v*_x_, and *v*_y_ as input. All models are symmetric such that mirror-symmetric inputs produce mirror-symmetric bout-type probabilities as output. All models pool data across *fed* and *starved* fish. **B-D.** Here we probe the strongest NN (red bar in **A**) with a synthetic prey object to see how bout-type probabilities depend on locations and properties of prey in the local environment. For simplicity, we show examples with a single synthetic prey at a single location (indicated with red circle), but this approach generalizes to arbitrary arrangements of potential prey. In **B**, we vary the size of the prey object (size in pixels; pixel side length = 13 µm) while holding its x-velocity and y-velocity constant. Similar to **4F** in main text, the “activations” of several bout-types are shown (panel 2), here for left and right versions of *e6*, *j2*, *p3*, *abort*, and *strike*. The full predicted bout-type probability distribution is evaluated at 3 indicated values for prey size (panels i-iii). With a prey object at this location, *p1-R* is always most likely, but *p1-R* probability increases as the object gets larger. In **C**, prey object x-velocity is varied from approaching the fish (panel i), to having zero x-velocity (panel ii), to moving away from the fish (panel iii). A speed of +1 pixel */* frame = +780 µm */* sec (away from the fish). With the object approaching the fish, high-energy exploring bouts are most likely. This is likely due to some residual coasting of the fish following its preceding swim bout, causing the relative x-velocity of an immobile object to be negative. With the object having zero or positive x-velocity, *p1-R* is most likely. In **D**, prey object y-velocity is varied from moving medially (panel i), to having zero y-velocity (panel ii), to moving laterally (panel iii). The y-component of velocity has a significantly nonlinear relationship to bout-type probability. At this location, the probability of *strike* is maximum as the object moves medially at ~780 µm */* sec (panel 2). With the object moving laterally, all rightward hunting bouts become much more likely, especially *p1-R*. **E.** Test-set F_1_-score (similar to **5H**) is shown for the best GLM model tested (purple bar in **A**). **F.** Test-set F_1_-score is shown for the best NN model tested (red bar in **A**). Prediction of all bout-types improves with the NN, especially for hunting bouts. **G.** Confusion matrix for best NN model (similar to **5I**). Notably, *strikes* are confused with *p1* bouts, likely due to the similar environmental conditions that elicit these bout-types but also the rarity of *strikes*.

**Figure S13:**
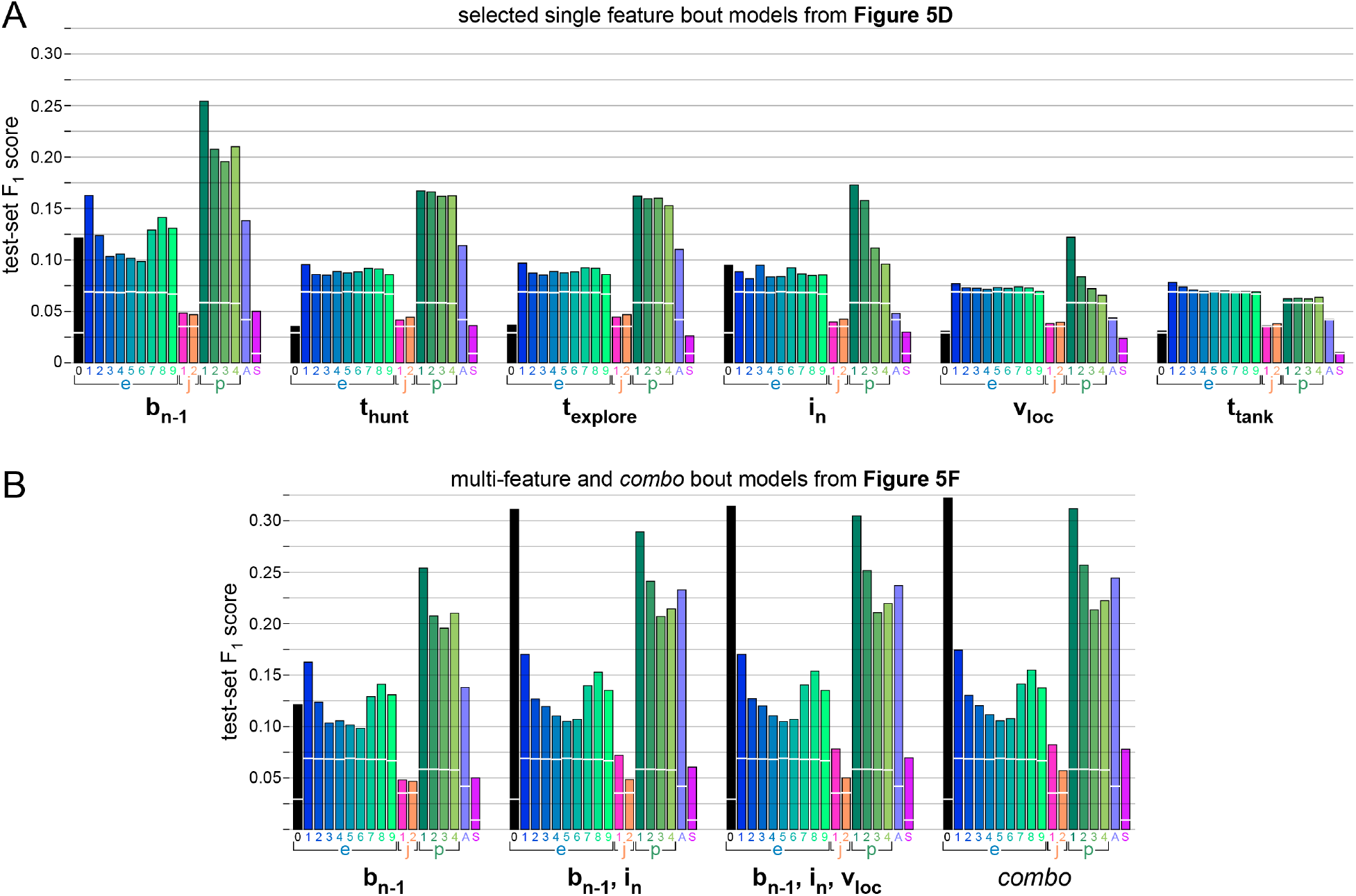
Comparing GLMs to predict bout-type on held-out data. **A.** Similar to **5H**, but for each selected single feature model described in **5D** of main text to predict bout-type (except *v*_size_, *v*_x_, *v*_y_). *Preceding bout-type* (*b_n_*_1_) is the best feature to predict every bout-type. **B.** Here we compare the *preceding bout-type* model (reproduced from **A**) to the best paired bout-type model ([*b_n_*_−1_, *i_n_*]), the best 3-feature bout-type model ([*b_n_*_−1_, *i_n_*, *v*_loc_]), and the *combo* bout-type model used in our *combo renewal process* simulations (Figure 6 in main text).

**Figure S14:**
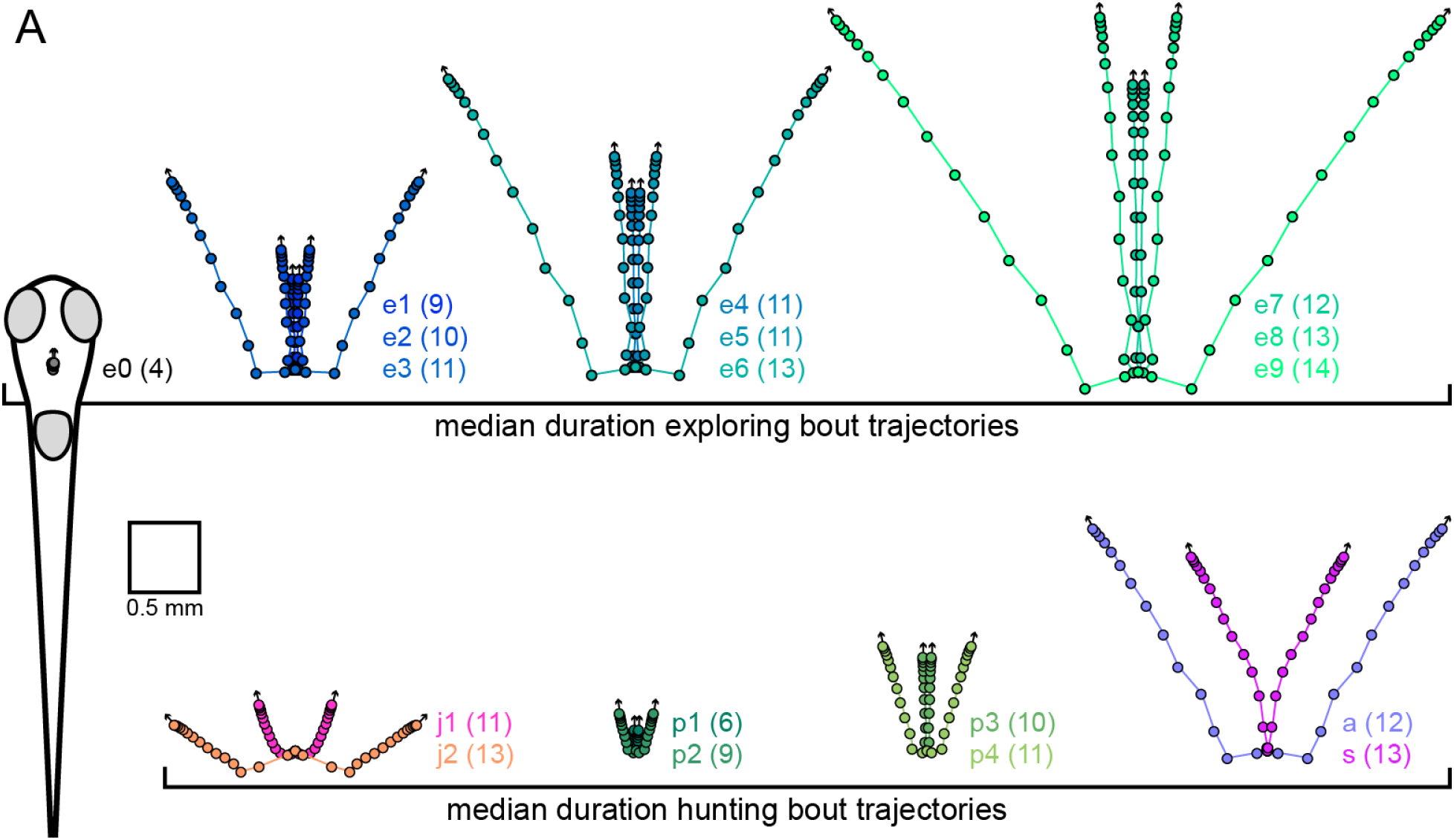
Bout-type trajectories used in *combo renewal process* simulations. **A.** Median duration (in number of frames) of each bout-type is shown in parentheses. The average trajectory for all median duration bouts of each bout-type are shown. Leftward and rightward versions of each bout-type are averaged to produce mirror symmetric trajectories. For each example trajectory, the fish is initially positioned as shown for the *e0* bouts, which have minimal displacement and heading angle change. We extend each trajectory to include the fish’s position through 3 additional image frames taken from the beginning of the following interbout interval (*i_n_*_+1_). This allows the fish to more realistically coast and decelerate following swim bouts in simulations. Circles indicate the fish head position during each frame, and the arrows extending from the terminal circle indicate the new heading direction at the end of each bout-type. These trajectories are used to move the fish through its virtual environment in the *combo renewal process* simulations described in Figure 6.

## Supplementary Video Legends

**Figure SV1: Data acquisition with the BEAST (animated).** This is an animation of the BEAST rig used to acquire data for this study. The larval zebrafish swims in an expansive arena (300 × 300 × 4 mm water volume) and is tracked with a camera moving automatically overhead. The tank rests on an air table to isolate the fish from the vibration of the moving gantry. The camera has an infrared filter and collects light from the array of IR-LEDs positioned on the air table. The diffusive screen embedded in the tank bottom scatters the infrared light and helps to create even lighting for behavioral imaging. A projector is used to project inwardly drifting gratings (not shown) onto the screen in the tank bottom to bring the fish to the center of the arena, where the camera waits to detect the arrival of the fish. Once at the center, the projector delivers a static color image of pebbles (pictured in the video).

**Figure SV2: Aligned behavioral video containing a hunt sequence.** This video shows the behavioral sequence described in **2J-K**. All behavioral video in this study is acquired at 60 frames per second, but this video is shown at half-speed (30 frames per second). The video contains 439 image frames (7.32 seconds in real time). The fish emits 2 exploring bouts, and then transitions into a 13-bout hunt sequence ending with a successful *strike* to capture a prey. The fish initiates the hunt by orienting toward a prey object with a high-|Δheading| leftward *J-turn* (*j2-L*). However, upon orienting toward this prey object, another prey arrives in the same location, and the fish appears to switch to a separate isolated prey target using a high-|Δheading|, high-|Δtail-shape| *pursuit* bout (*p4-L*). The fish then pursues this moving prey object, closing the distance to the target and aligning its heading direction with the heading direction of the target. During prey capture, the prey can be seen entering the mouth of the fish and then moving around inside the mouth before it is swallowed. Swallowing of the prey coincides with emission of an *e0* bout. These subtle movements highlight a challenge in parsing larval zebrafish behavior into bouts and interbouts since swallowing behavior and other orofacial and pectoral fin movements are near our threshold for bout detection. In the right panel of the video, the 20 tail vectors used to define tail-shape are shown in each image frame, and the vergence angle of each eye is indicated by coloring the outline of each eye with the “cool” colormap from Matlab, ranging from blue (diverged) to magenta (converged). Prey objects are extracted from the movie for display with image processing routines similar to those used to encode environmental state preceding each swim bout.

**Figure SV3: Animation of a simulated behavioral trajectory** The behavioral trajectory shown in **6A** is reproduced here in an animation. This excerpt was originally 411 frames (6.85 seconds of simulated time), but is slowed down 3x for display in this video. This behavior is generated by sampling from our *combo renewal process* model (see Methods). In this sequence, the virtual fish exits exploring mode to hunt a moving prey object, ending this virtual hunt with a *strike* toward the prey. This example captures many salient features of naturalistic larval zebrafish behavior observed in our study. That being said, we searched our simulated data to locate a behavioral sequence for display, and this example is not necessarily representative of an average simulated hunt. Also, when the fish *strikes* at the virtual prey object in this movie, we remove that prey from the video for display purposes. However, in our simulations, the environment runs in open-loop and prey are not affected by the behavior of the virtual fish.

**Figure SV4: Visualizing action selection in *combo renewal process* simulations** Here we show a full 2-minute behavioral trial simulated from our *combo renewal process* model. This video is slowed down 4x relative to simulated time. The behavioral sequence shown in **6A** is taken from this simulated trial (beginning with bout *b*_161_ at 6:14 in the video). Fish head position (colored circle) and heading direction (rightward) are held constant, and the virtual environment translates and rotates as the fish moves during swim bouts. Virtual prey are shown as black circles and the corresponding representation of prey locations with the *v*_loc_ feature is shown (upper right panel). For simulation (see **6A**, Methods, Technical Appendix for more detail), an interbout interval duration is randomly sampled from a probability distribution generated from the *combo* interbout model (bottom right in video). Sampled interbout interval durations are indicated with a red bar, and the distribution from which it was sampled is shown in black. A gray bar moves rightward to indicate the passage of time during interbout intervals. As time passes, we update the bout-type probability distribution generated from our *combo* bout model (bottom left of video), evaluated for the currently displayed value of the ongoing interbout interval (this is the preceding interbout interval on which the next bout-type partially depends). In this way, the change in bout-type probabilities with increasing interbout interval can be observed. Once the sampled time for the interbout interval has elapsed, a bout-type is randomly sampled from the bout-type probability distribution (indicated with a star). The virtual fish then moves through its environment along the designated trajectory for that bout-type (see S14).

## Technical Appendix

This appendix describes the details of our mathematical notation, feature encoding, construction of the marked renewal process model, procedures for fitting the model, methods for obtaining posterior credible intervals of key parameters, and methods for hyperparameter selection.

### Notation

The data consists of many bout sequences, as described above. Let 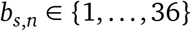 denote the bout type of the *n*-th bout in the *s*-th sequence, and let 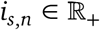 denote the preceding interbout interval. Let *N_s_*denote the number of bouts in the *s*-th sequence and *S* denote the total number of sequences. The set 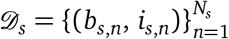 is the sequence of bouts and interbout intervals in the *s*-th sequence, and 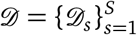 is the set of all sequences’ data.

The features are functions of the preceding bouts and interbout intervals and of the current environmental input. Let 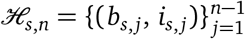 denote the history of bouts and interbout intervals preceding the *n*-th bout in sequence *s*.^1^ These histories will form the input to the interbout interval and bout type GLMs. Let 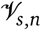 denote the set of environmental inputs to the *n*-th bout in the *s*-th sequence, and let 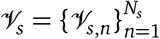 denote the sequence of environmental inputs for each bout in the sequence.

Feature functions 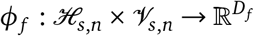 map behavioral history and environmental inputs to a vector representation of length *D_f_* that can be input to a GLM. Here, *f* denotes the type of feature, like previous bout type or preceding interval, and each feature type may have its own dimensionality. The feature functions we consider only use one of the inputs at a time, though more complex feature functions could be used in future work. Next we specify how these feature functions are computed.

### Feature Encoding

For clarity, we omit the subscript-*s* from the following definitions as the feature functions are not sequence-dependent. The first feature is a constant bias, 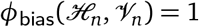. This allows for baseline probabilities of different bout types and a baseline mean and dispersion of the intervals.

Discrete features like the preceding bout type are represented with a one-hot encoding. If the preceding bout type is *b_n−_*_1_ = *k*, we have

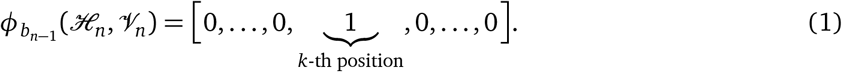

Scalar features like *i_n_*, *t*_hunt_, *t*_explore_, and *t*_tank_ are encoded with a set of basis functions. Let 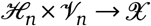 denote a generic function that maps behavioral history and environmental input to a scalar value in the set 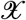. For example, the preceding interval is a function of the behavioral history that outputs a non-negative scalar value. We use a data-driven approach to determine a set of basis functions for representing these scalar values in a way that allows the GLMs to learn nonlinear dependencies. First, we compute the empirical cumulative distribution function (CDF) 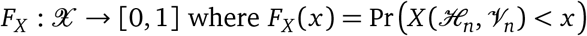. We approximate this function by computing the histogram of *X* values on a fine grid (typically with 100 equispaced bins). To construct a basis that has higher precision around the most common values, we map 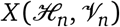 through the CDF to obtain a point [0, 1], and then we evaluate a set of *J* orthogonal basis functions at that point to obtain a feature vector. Formally, we have,

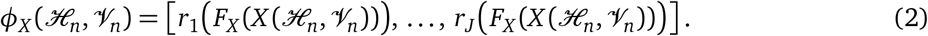

The basis functions 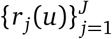 are obtained by orthogonalizing a set of radial basis functions 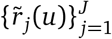, where,

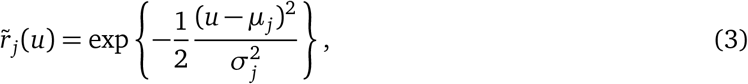

for *u ∈* [0, 1]. We evenly space means at values 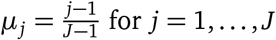 and set the standard deviation to 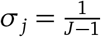. Finally, we perform a QR factorization to obtain the orthonormal basis on the interval [0, 1]. As discussed below, each feature type has a different value of *J*, as chosen via an empirical Bayes procedure. Figure S9 shows the bases computed on a dense grid of points in 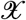 for a variety of features.

Finally, we have a set of environmental features including the location, size, *x*-velocity, and *y*-velocity of the paramecia in the fish’s field of view. Each of these features is a scalar value for each paramecium in the field of view. We summarize the activity of all paramecia by summing the feature values for all paramecia within a spatial bin. For small enough bins, this representation entails little loss of information, but future work could consider more sophisticated feature functions. The spatial bins are constructed by dividing the angular and radial dimensions of the fish’s field of view into bins. A bin that spans *Δθ* radians in the angular dimension and covers the interval [*r*, *r* +*Δr*) in the radial dimension has area 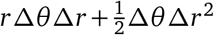. This area grows with the distance from the fish, *r*. To account for this difference in area, we normalize the summed feature values by the area to get a “feature density” within each bin. As with the temporal bases, we choose the number of angular and radial bins via the empirical Bayes procedure described below.

### Constructing the Marked Renewal Process

The marked renewal process specifies a conditional probability density over a sequence of bouts and interbouts given model weights 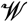. By the chain rule, this density factorizes into a product of conditional probabilities of each interval and bout type given all preceding bouts and intervals, and, by assumption, the *n*-th bout depends only on the current environment. Formally,

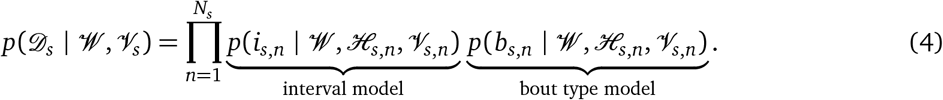

We assume that all sequences share the same set of weights and are conditionally independent given the weights so that 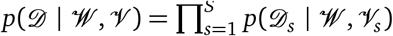.^2^

We use generalized linear models (GLMs) for both the intervals and the bout types. In a GLM, we first compute a weighted combination of the features via the “activation functions.” The output of the activation functions are then passed through a fixed nonlinear function to obtain the parameters of the conditional distribution on intervals or bout types, as appropriate. Typically, there is a separate activation function, and hence a separate set of weights, for each parameter in the final conditional distribution.

#### Interbout Interval GLMs

First consider a Poisson interval model parameterized by its mean 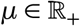. The mean is obtained from the bout history and environmental input as follows. Let 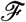 denote the set of features input to the GLM. The activation function for the mean is given by,

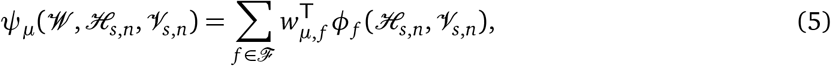

where 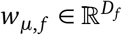 is a weight vector in the set of weights 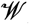, and it is specific to this parameter (*µ*) and feature (*f*). It is the same dimensionality as the output of the feature function *φ_f_*. Thus, the output of the activation function is a real-valued scalar. Finally, the output of the activation function is passed through an inverse link function (here, an exponential function) to obtain the mean,

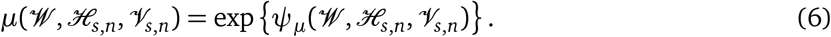

Multi-parameter conditional distributions, like the negative binomial distribution parameterized by a mean and dispersion, require two separate activation functions and sets of weights. Table 1 lists the various forms of generalized linear models we used to model interbout intervals.

**Table 1:**
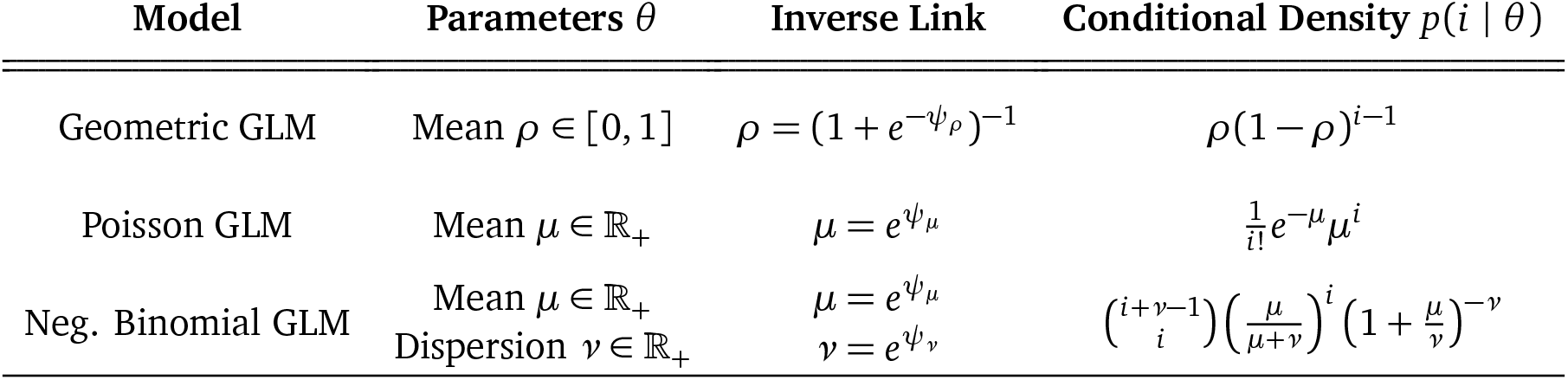
Conditional distributions for interbout intervals *i*, measured in number of elapsed frames between bouts.

| Model | Parameters $\theta$ | Inverse Link | Conditional Density $p(i \vert \theta)$ |
| --- | --- | --- | --- |
| Geometric GLM | Mean $\rho \in [0, 1]$ | $\rho = (1 + e^{-\psi_\rho})^{-1}$ | $\rho(1 - \rho)^{i-1}$ |
| Poisson GLM | Mean $\mu \in \mathbb{R}_+$ | $\mu = e^{\psi_\mu}$ | $\frac{1}{i!} e^{-\mu} \mu^i$ |
| Neg. Binomial GLM | Mean $\mu \in \mathbb{R}_+$<br>Dispersion $\nu \in \mathbb{R}_+$ | $\mu = e^{\psi_\mu}$<br>$\nu = e^{\psi_\nu}$ | $\binom{i+\nu-1}{i} \left(\frac{\mu}{\mu+\nu}\right)^i \left(1 + \frac{\mu}{\nu}\right)^{-\nu}$ |

#### Bout Type GLMs

We modeled the conditional distribution of the next bout type in a similar fashion. The bout type follows a categorical distribution parameterized by *π ∈ Δ_K_*, a non-negative vector of length *K* that sums to one; i.e. a probability distribution over the *K* = 36 bout types. We modeled this probability vector with a GLM with a softmax inverse link function,

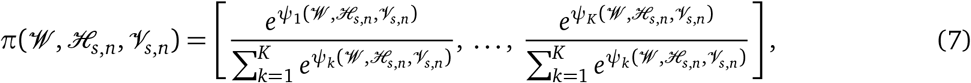

where 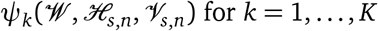 is an activation function that specifies the relative likelihood of the next bout type being of type *k*.

#### Weight Sharing

We assessed between-group differences and lateral biases by tying the weights for some features and comparing model performance. For example, to assess the lateral symmetry we compared the standard bout-type GLM described above to a “symmetric” GLM in which the weights for left and right bout types are constrained to be the same. More specifically, if *k* and *k^!^* correspond to the left and right versions of a bout type, like a particular J-turn, we constrain 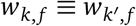 for symmetric features *f*.

Enforcing left/right symmetry is slightly more challenging for the weights of the preceding bout type feature. In that case, we can think of the collection of weights as a *K × K* “transition” matrix. If the bout types are sorted so that the first half correspond to left and the second half correspond to right, we tie the weights of the upper-left and lower-right quadrants (i.e. the ipsilateral transition weights), as well as those in the upper-right and lower-left quadrants (i.e. the contralateral transition weights). Likewise, we enforce symmetry in the environmental input weights by tying the weights of the each hemisphere of the field of view and its ipsi- or contralateral bout types.

#### Prior Distributions

We introduce a Gaussian prior on the model weights in order to regularize against overfitting, particularly with high-dimensional features. For each parameter *θ* and feature *f* we have,

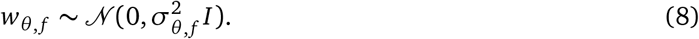

We learn the variance 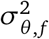 via the empirical Bayes procedure described below. We considered Laplace priors, which are akin to 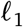 regularization, as well, but initial explorations showed minimal differences from the Gaussian distribution.

### Fitting the Model

We fit the model with maximum *a posteriori* (MAP) estimation. We computed the log joint probability density,

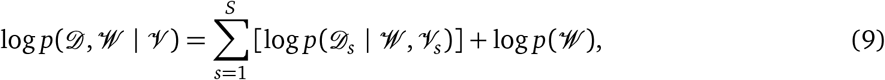

where the first term is the log likelihood from (4) and the second is the prior from (8). We used Autograd (https://github.com/HIPS/autograd/) to compute its gradients with respect to the model weights 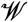. We optimized the log joint probability with Newton’s method, using the conjugate gradient method to solve the linear system in the Hessian matrix 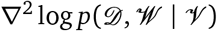, as implemented in SciPy.

Once the MAP estimate of the weights 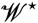 was obtained, we computed a Laplace approximation to the posterior distribution,

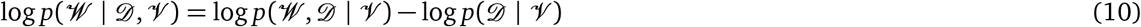

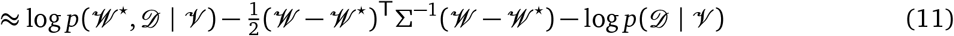

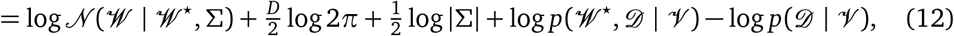

where the inverse covariance matrix is the negative Hessian at the mode,

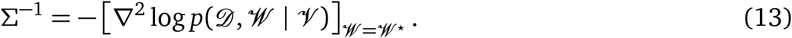

(Note that we have committed a slight abuse of notation by treating 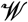 as a vector rather than a set. To be precise, take 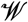 to be the concatenation of the vectorized weights.)

#### Posterior Credible Intervals

We obtain approximate posterior credible intervals from the posterior covariance matrix *Σ*. Consider a particular parameter *θ* and feature *f*. Let 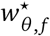 denote its posterior mean and *Σ_θ_* _, *f*_ denote its posterior covariance. (The covariance of this single parameter and feature’s weights is just one diagonal block of the complete covariance matrix *Σ*.) The 95% posterior credible interval for the *d*-the entry in the weight vector is approximately 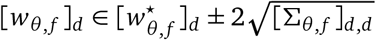. For scalar features represented by basis function expansions, we obtain the credible intervals like those shown in Figure 3F by first constructing a matrix of basis function evaluations 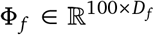 where the *t*-th row is *φ_f_* evaluated at the *t*-th percentile of the feature values. Then we compute the posterior credible interval in feature space as. 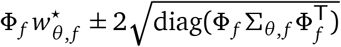

### Empirical Bayes Hyperparameter Selection

Since the posterior density (12) integrates to one over 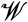, the Laplace approximation also yields an approximation to the marginal log likelihood,

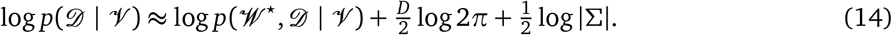

The marginal likelihood is the probability of the data, integrating out the weights under their prior distribution. As seen in (14), it increases with the log joint probability density at the mode 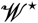, but it decreases as the curvature at the mode grows and the posterior covariance *Σ* shrinks to zero. That is, the marginal likelihood balances posterior probability density around the mode with posterior uncertainty. A model with high marginal likelihood should assign high probability to the data over a large set of weights. This balance makes the marginal likelihood a natural measure of model fitness.

We select the hyperparameters—specifically, the prior variances 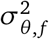 and the number of basis functions or spatial bins *J* —by comparing the marginal likelihood estimates over a grid of values *for each single-feature model*. For each feature, we scan over a grid of hyperparameter values, and for each hyperparameter setting we fit a GLM with only that feature and the bias. We approximate the marginal likelihood using (14) and select the hyperparameter setting that achieves the maximum value over the grid. We then use these hyperparameter settings on composite models that combine multiple features. While the optimal single-feature hyperparameters are not necessarily optimal for the full model, we expect the selection to be biased toward over-parameterized models with more basis functions and larger prior variances. Erring in this direction is more conservative, in that it allows the full models greater flexibility, even if it is not necessary.

### Neural Network Models of Environment Dependence

We also explored artifical neural network models to learn more complicated environment dependencies in the bout type GLMs. For these models, we treated *φ_v*_*, where *\** denotes one of loc, size, x, or y, as the input to a multilayer feedforward network whose last layer is a softmax, as above, to output a probability distribution over the *K* = 36 bout types. Again, we use weight sharing to enforce lateral symmetry in the input-output function, but here the weights must be shared within each layer. Since the training function is no longer convex, we instead train the model with the Adam^90^ optimizer using mini-batches of randomly selected sequences. We use cross-validated log likelihood to determine the number of layers and hidden units. The Laplace approximation is not appropriate for this multimodal log probability, so we instead compare models on the basis of the log likelihood they assign to test data.

## Footnotes

1 For bout models, the history also includes *i_s_*,*_n_*, the interval preceding the *n*-th bout.

2 The first bout/interbout interval (*b_s_*_,1_, *i_s_*_,1_) require some care as, at that point, there is no history to condition on. Likewise, the last interval is truncated if tracking is lost. We do not explicitly model these effects; instead, we assume the sequences are stationary and let the initial distribution be the empirical distribution of bouts and intervals, and we drop the last interval from our model. Since the vast majority of bouts are not at the start or end of the sequence, these assumptions have little effect on our inferred model parameters.

